# Structural basis of organic cation transporter-3 inhibition

**DOI:** 10.1101/2022.07.14.499921

**Authors:** Basavraj Khanppnavar, Julian Maier, Freja Herborg, Ralph Gradisch, Erika Lazzarin, Dino Luethi, Jae-Won Yang, Chao Qi, Marion Holy, Kathrin Jäntsch, Oliver Kudlacek, Klaus Schicker, Thomas Werge, Ulrik Gether, Thomas Stockner, Volodymyr M. Korkhov, Harald H. Sitte

## Abstract

Organic cation transporters (OCTs) facilitate the translocation of catecholamines and xenobiotics across the plasma membrane in various tissues throughout the human body. OCT3 plays a key role in low-affinity, high-capacity uptake of monoamines in most tissues including heart, brain and liver. Its deregulation plays a role in diseases. Despite its importance, the structural basis of OCT3 function and its inhibition has remained enigmatic. Here we describe the cryo-EM structure of human OCT3 at 3.2 Å resolution. Structures of OCT3 bound to two inhibitors, corticosterone and decynium-22, define the ligand binding pocket and reveal common features of major facilitator transporter inhibitors. In addition, we relate the functional characteristics of an extensive collection of previously uncharacterized human genetic variants to structural features, thereby providing a basis for understanding the impact of OCT3 polymorphisms.

## Introduction

Organic cation transporters (OCTs; Fig. 1a) are low-affinity, high-capacity transporters^1^ which act complementary to high-affinity, low-capacity neurotransmitter:sodium symporters (solute carrier family 6, SLC6)^2^ in regulating and maintaining the extracellular equilibrium of monoamine neurotransmitters^3^. Norepinephrine, dopamine and serotonin belong into this family; interaction with their cognate receptors and subsequent signaling events are essential for physiological function and play multiple roles in a variety of pathologies^4,5^. In addition, OCTs are responsible for cellular uptake of cationic xenobiotics in various tissues^1^.

**Figure 1.**
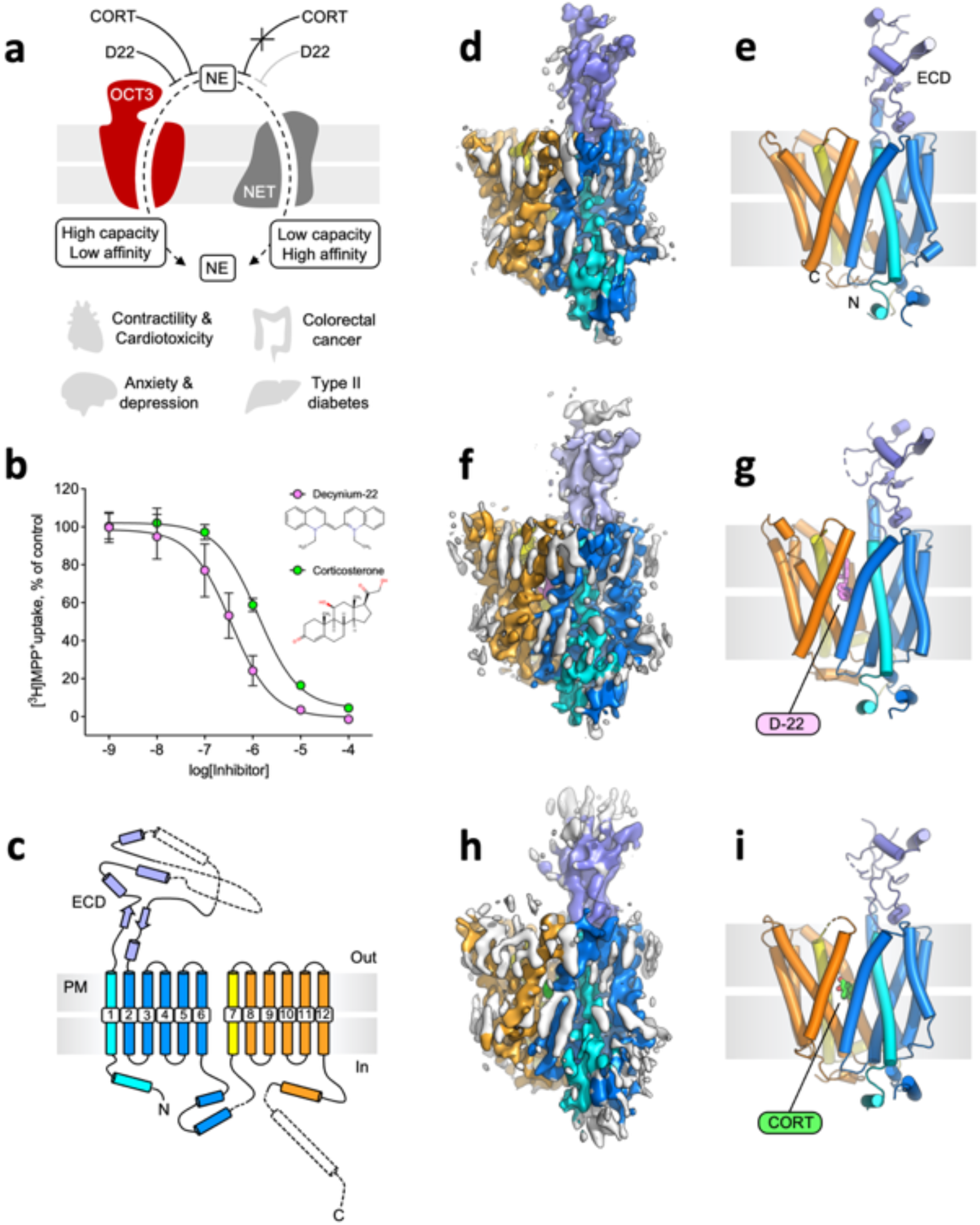
Structure and function of OCT3. **a**, Schematic representation of OCT3; key features of the transporter are illustrated in the panel. **b**, OCT3 transport inhibition by decynium-22 (D22) and corticosterone (CORT). The values correspond to mean ± SD; n = 3, in triplicate. **c**, A scheme depicting the topology and the secondary structure elements of OCT3. **d-e**, The cryo-EM map (d) and model of OCT3 in nanodiscs at 3.2 Å resolution. The colors of the protein correspond to those in c; annular lipids are colored grey. **f-g**, Same as d-e, for OCT3-D22 complex at 3.6 Å resolution (D22 colored violet). **h-i**, Same as f-g, for OCT3-CORT complex at 3.7 Å resolution (CORT colored green).

Organic cation transporter 3 (OCT3, SLC22A3) was identified as a corticosterone-sensitive catecholamine transporter in 1998 and belongs to the SLC22 family of the major facilitator superfamily (MFS) of transporters^6^. OCT3 is more widely expressed than OCT1 and OCT2, which are primarily located in liver and kidney, respectively^7^. Due to the effect of OCTs on the pharmacokinetic fate of therapeutically relevant drugs, FDA and EMA recommend to screen compounds for possible interaction with OCTs^8^. A number of single nucleotide polymorphisms (genetic variants) have been identified in OCT3; in some instances the missense mutations have been linked to specific functional properties of the transporter and associations identified between genetic variants and cardiovascular disease^9,10^, type 2 diabetes^11,12^ and cancer^13^.

Recent findings reinforce the role OCT3 of in various (patho-)physiological processes. Norepinephrine uptake in cardiomyocytes is necessary for cardiac contractility, which is mainly mediated by OCT3^14^. OCT3 also mediates the uptake of doxorubicin into the myocardium and thus contributes to its dose-limiting toxicity^15^. OCT3 polymorphisms and reduced hepatic OCT3 expression, caused by cholestasis and liver fibrosis, lead to reduced metformin uptake, thereby impacting its therapeutic action through changed pharmacokinetics^7,16,17^. In addition, loss of OCT3 is associated with progression of liver fibrosis^18^ and hepatocellular carcinoma^19^. Colorectal cancer cells overexpress OCT3 which renders them susceptible to oxaliplatin-induced cytotoxicity^20^. Finally, inhibition of OCT3 allows for raising extracellular levels of serotonin and norepinephrine in the brain and may thus present an alternative to currently approved antidepressants in the treatment of depressive disorders^21,22^.

Potent and selective inhibition of OCT3 may have immediate medical implications. However, specific ligands are scarce and progress is hampered by the lack of structural information. Prototypical ligands of OCT3 are decynium-22 (D22) and the OCT3-selective steroid hormone corticosterone (CORT). We set out to determine the structure of OCT3 alone, as well as in complexes with its two characteristic inhibitors. In addition, we delineated the structure-function relationship of OCT3 by examining how genetic variants of OCT3, which were discovered by exome-sequencing in a cohort of 17,339 subjects, affect transport activity.

## Results

### Structure determination of human OCT3

We first set out to determine the structure of OCT3 using cryo-EM and single particle analysis. We purified and reconstituted OCT3 into nanodiscs (MSP1D1 filled with brain polar lipids; Supplementary Fig. 1a-b) and subjected the sample to extensive cryo-EM data collection and image processing (Supplementary Fig. 1c and 2, Supplementary Table 1). We obtained a 3D reconstruction of OCT3 (apo-state) at 3.2 Å resolution (Figure 1d-e, Supplementary Fig. 2, 5a-b and 6) and completed the model by combining the experimentally determined structure with an *in silico* model generated by AlphaFold (detailed in Materials and Methods, Supplementary Fig. 6 and 7f).

The structure revealed a classical MFS fold for OCT3, with the transporter composed of twelve TM helices (TM1-12) in an outward-facing conformation. The translocation pathway is located at the interface of the two 2-fold pseudo-symmetrically-related transmembrane domains consisting of TM1-6 and TM7-12. The substrate binding site is located in the center of the transporter between the two domains, halfway through the membrane. The structure features a prominent, partially resolved density of the extended extracellular loop 1 or ectodomain (ECD), which is adjacent to the outward-open translocation pathway (Fig. 1c-d). Upon expression in human embryonic kidney-293 cells, human OCT3 transported its substrate MPP^+^ (1-methyl-4-phenylpyridinium) and showed inhibition by two key molecules, D22 and CORT (Fig. 1b).

### Structures of D22- and corticosterone-bound OCT3

We incubated purified and reconstituted OCT3 with saturating concentrations of D22 and CORT (1 mM, Fig 1b). We subsequently determined the structures of OCT3 in D22- and in CORT-bound states at 3.6 Å and 3.7 Å resolution, respectively (Fig. 1f-i, Supplementary Fig. 3, 4 and 5c-f). A comparison of the two inhibitor-bound states with the apo-state of OCT3 clearly showed that the compounds are readily accommodated by the protein in its outward-facing conformation with minimal conformational changes (Fig. 2, Supplementary Fig. 8). The root mean square deviation (RMSD) between all atoms of the apo-state and each of the inhibitor-bound states is 0.01 Å (apo vs D22) and 0.69 Å (apo vs CORT). CORT placement was assisted by molecular dynamics (MD) simulations, as detailed in “Materials and Methods” (Supplementary Fig. 7, 8 and 9). The cryo-EM structures of OCT3 and the full-length models of the loops, completed with AlphaFold, were further validated using MD simulations. The trajectories showed stable structures for apo-OCT3 as well as for the D22- and CORT-bound structures (Supplementary Fig. 7 and 9). Addition of the AlphaFold-based missing parts of the extracellular and intracellular domains increased the stability of OCT3 secondary structure and reduced the deviation from the starting structures (lower RMSD and root mean square fluctuation (RMSF), Supplementary Fig. 7).

**Figure 2.**
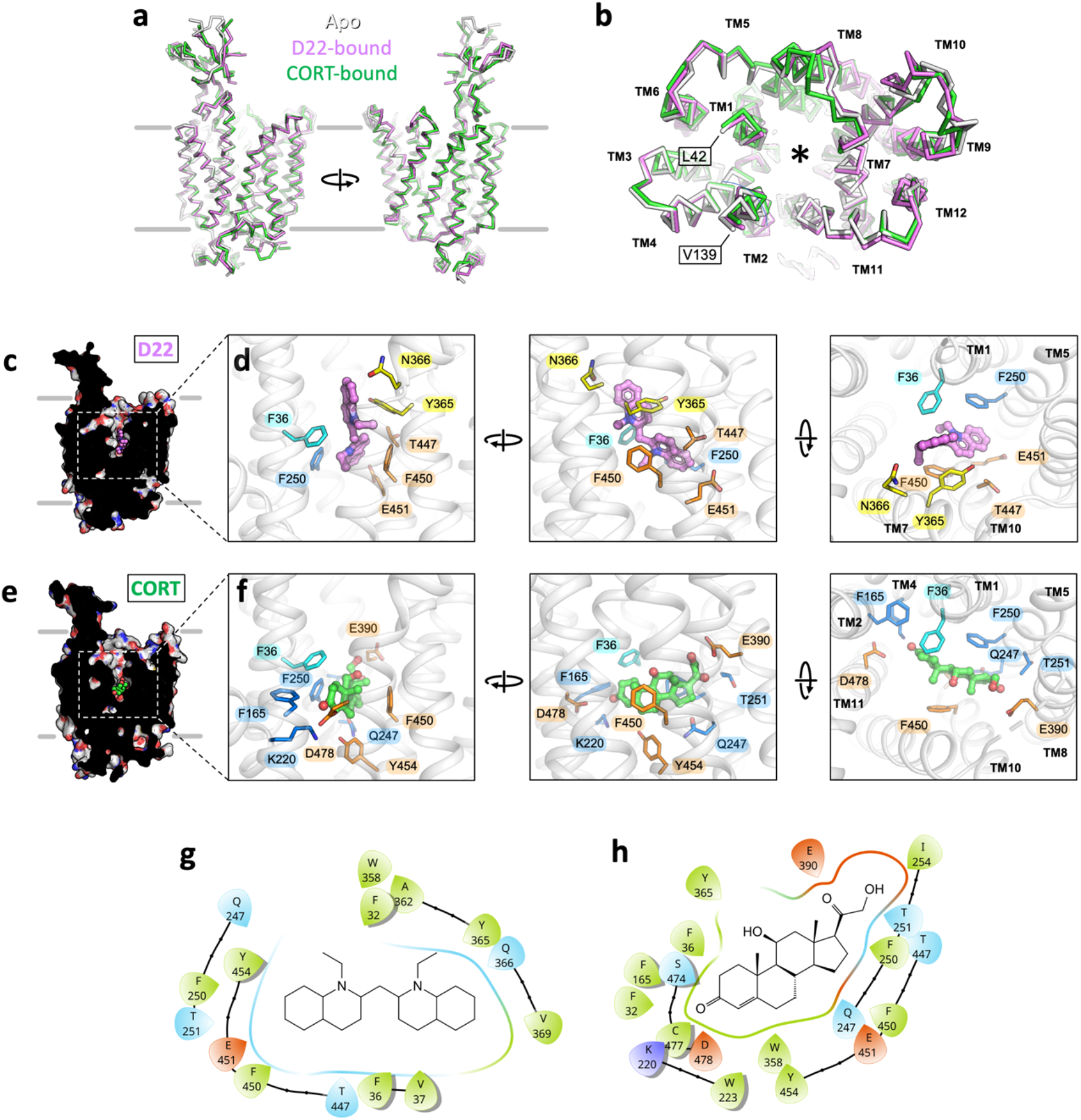
Comparison of the apo-state and the ligand-bound states of OCT3. **a**, Structural alignment of the three indicated states of OCT3 shows an overall high degree of similarity. **b**, same as in a, view from the extracellular space. The positions of each transmembrane (TM) helix are indicated. The residues flanking the extended extracellular loop 1 (ECL1) are indicated with boxes (L34 and V139). The translocation pathway / ligand binding site is indicated by an asterisk. **c**, A sliced view of D22, showing the inhibitor buried deep in the substrate translocation pathway. **d**, The expanded views of the D22 binding site in different orientations (*left* the same orientation as the one in c), indicating the residues within 4 Å distance of the inhibitor. **e-f**, Same as c-d, for OCT3-CORT. The TM domains containing the residues in close proximity to the ligand are indicated in the right-most panel (d and f). **g-h**, 2D interaction plot showing the residues interacting with D22 and CORT.

Both compounds occupy the substrate translocation pathway resulting in its steric blockage. Interestingly, the binding sites for D22 and CORT partially overlap, but the orientations of the compounds within the pocket are distinct. The cationic D22 is bound to OCT3 in a pose oriented perpendicular to the membrane plane (Fig. 2d). D22 binds to only one side of the outer vestibule, making several interactions with TM7 and TM11 (Fig. 2g). In contrast to D22, CORT is positioned roughly parallel to the plane of the lipid bilayer (Fig. 2f) and makes more extensive contacts within the binding pocket, with 11 residues (Fig. 2h).

Although a number of residues are shared between the binding sites of CORT and D22, the mode of binding of these two compounds differs substantially (Fig. 2c-h). Furthermore, in the presence of CORT, the OCT3 density map features several additional elements located at the entrance to the outward-open binding pocket (Supplementary Fig. 10a). We interpret this as a low-affinity site that may accept weakly bound CORT molecules in alternate poses (Supplementary Fig. 10b). The poses of D22 and CORT suggest that the mechanism of transport inhibition by these two compounds may differ. The tightly bound D22, positioned perpendicular to the membrane plane within the binding site, may prevent outward to inward conformational changes in OCT3. In contrast, CORT occupies a larger footprint within the binding site. It may also prevent outward-to-inward rearrangements of the transporter by occupying multiple non-specific binding sites and thus “clogging” the translocation pathway.

### Molecular basis of OCT3 ligand specificity

CORT is selective for OCT3^6,23,24^. In contrast, D22 binds to and inhibits all OCTs (Fig. 3e), showing only slightly higher affinity for OCT3. However, residues interacting with D22 and CORT in OCT3 are largely conserved among the three OCTs (Fig. 3f). The ligand binding sites differ in only six residues: F36 (TM1), F250 (TM5), I254 (TM5), F450 (TM12), E451 (TM12) and Y454 (TM12). These six residues correspond to C36, F244, L248, I446, Q447, Y454 in OCT1 and to Y37, Y245, L249, Y447, E448, C451 in OCT2 (Fig. 3f). We compared the OCT3 structures to the homology models of OCT1 and OCT2 generated by AlphaFold^25^ (Fig. 3g). The OCT1 substrate binding pocket lacks negatively charged residues, compared to OCT2 and 3. The tyrosine residues in OCT2 (Y245, Y447) that correspond to F250 and F450 in OCT3 increase the hydrophilicity of the substrate binding pocket and provide two additional hydrogen bond donors.

**Figure 3.**
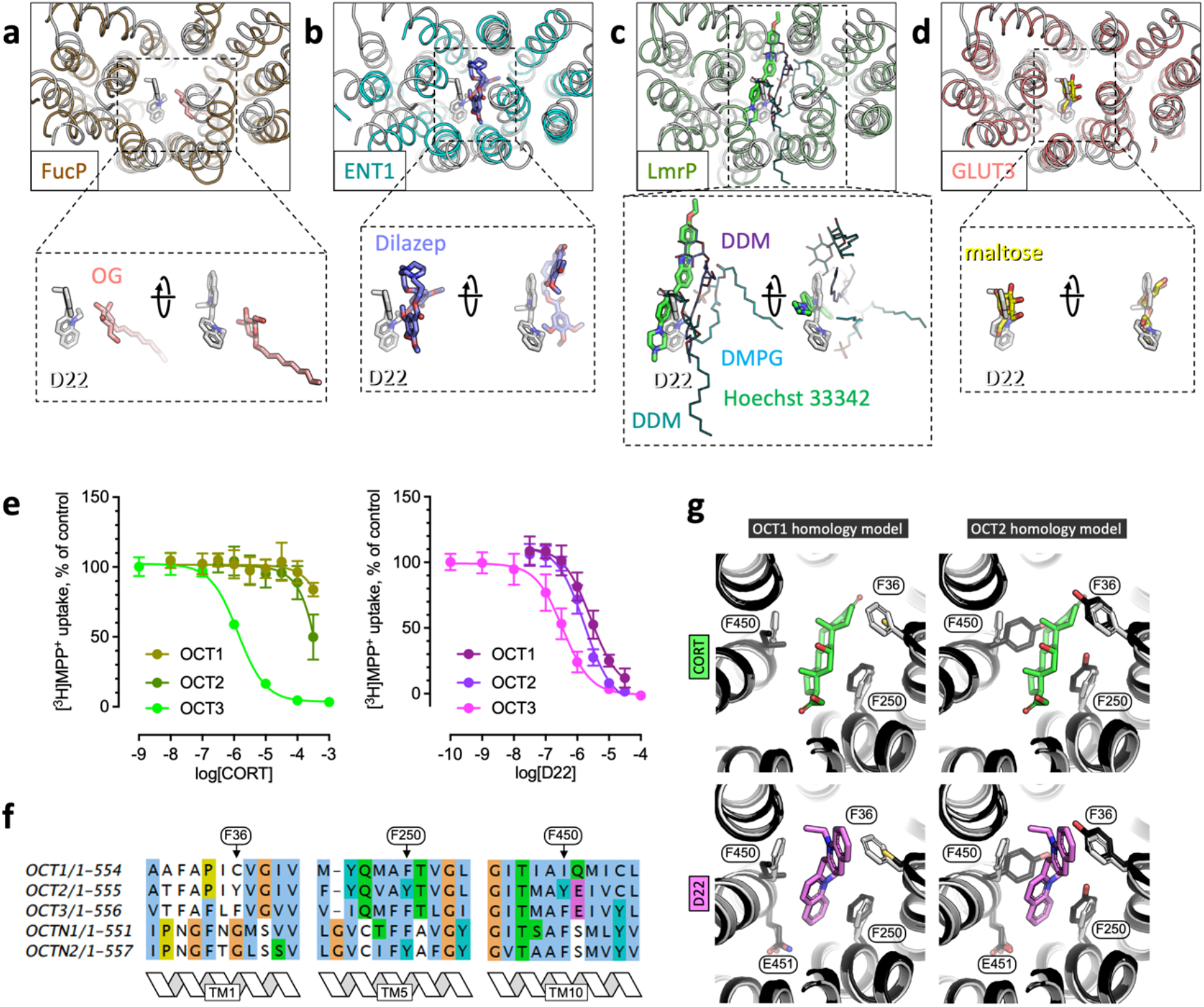
Molecular basis of OCT3 ligand specificity. **a-d**, Comparison between OCT-D22 and four different MFS transporter structures in outward-facing, ligand bound states, including FucP (PDB ID: 3o7q), ENT1 (PDB ID: 6ob7), LmrP (PDB ID: 6t1z) and GLUT3 (PDB ID: 4zwc). The dotted lines/boxes show the zoomed in views of the isolated ligands (*left*, same orientation as in the main panel; *right* – rotated ∼90°). **e**, Inhibition of OCT1, 2 and 3 transport by D22 and CORT (mean ± SD; n = 3). **f**, Sequence alignment of OCT1-3, OCTN1 and OCTN3, indicating the positions of the key OCT3 residues involved in ligand binding (and varied among the homologues): F36, F250 and F450. **g**, Comparison of the CORT- and D22-bound states in the experimentally determined OCT3 structures (white) with the OCT1 and OCT2 homology models (black).

### Negatively charged residues line the translocation pathway

The movement of organic cations across the plasma membrane earned OCTs their name^26^. The prerequisite for efficient cation transport is the presence of a regulated anionic surface that attracts the cationic ligand on one side of the membrane, induces a conformational change of the transporter and leads to the release of the compound on the other side of the membrane in one productive transport cycle. The structure of OCT3 features various negatively charged residues lining the substrate translocation pathway: D155, E232, D382, E390, E451, E459 and D478. Several of these residues are in proximity of the OCT3-bound D22 and CORT molecules (Supplementary Fig. 11a). A comparison of the OCT3 structure with the homology models of OCT1 and OCT2, the two closest homologues of OCT3 in the SLC22 family, as well as with the organic anion transporters (OATs, also members of SLC22 family of transporters) illustrates the negative and positive electrostatic potential in OCT and OAT translocation pathways, respectively (Supplementary Fig. 11). The electrostatic properties of these transporters appear to be consistent with their functional annotation as anion or cation transporters. The differences in charge distribution within the substrate binding pockets of OCTs and OATs are consistent with their substrate preference (Supplementary Fig 11).

### Comparison of OCT3 ligand binding to other MFS transporters

Despite the availability of several MFS transporter structures, only few have been determined in states bound to their small molecule substrates or to inhibitors^27-32^. A comparison of OCT3-bound D22 and OCT3-bound CORT to other small molecule-bound transporters (Fig. 3a-d) captured in an outward-facing state shows that the inhibitors of these transporters utilize a common, simple and effective mechanism of inhibition: binding to the substrate binding site in connection with inhibition of conformational changes essential for substrate translocation. The region of the transporter involved in these inhibitory interactions involves residues in TM1, TM2, TM5, TM7, TM8 and TM11, which are exposed to the substrate translocation path (Fig. 3a-d). As noted above, in OCT3 the orientation of D22 perpendicular to the membrane plane may not only block the translocation pathway, but may also prevent an outward-to-inward rearrangement of the OCT3 conformation. It is possible that the effective inhibitors of the other MFS transporters share this feature with D22, occupying a position that blocks the translocation pathway and constrains the conformational flexibility of the protein. This may have important implications for rational design of MFS transporter inhibitors.

### Lateral access into the substrate translocation pathway

The outward-open state of the lipid-reconstituted OCT3 is surrounded by several lipid densities (Fig. 1d). The protein structure features a V-shaped lateral opening at the interface between the two halves of the protein (“V-site”; Supplementary Fig. 12). Several conserved lipid densities are present at this site, indicating the margins of the lipid bilayer (Supplementary Fig. 13). We presently cannot unambiguously assign the identity of the lipids in this region (brain polar lipids, which include cholesterol and phospholipids, were used for nanodisc reconstitution). This V-site may serve as an access pathway for hydrophobic molecules that diffuse into the OCT3 translocation pathway. Although similar features are present in other MFS transporters^27^, the structure of OCT3 in a nanodisc allows us to visualize the lipid densities at this lateral opening (Supplementary Fig. 12).

### Genetic variants of OCT3

We investigated the occurrence of coding genetic variants in a large exome sequencing dataset from the iPSYCH2012 cohort^33^. This dataset includes 4885 healthy controls and 12454 patients diagnosed with at least one of five major psychiatric diseases: ADHD, autism-spectrum disorders, bipolar disorder, depression or schizophrenia. In total, we identified 58 different coding variants in 402 individuals in the combined cohort of cases and controls (Supplementary Table S4). These included 27 novel and 31 previously reported variants according to the Genome Aggregation Database database^34^. We then performed carrier-based association analyses to compare the carrier frequency of coding variants among control subjects and patients. Remarkably, we found a significant enrichment of coding variants in control subjects with a 1.29 fold higher carrier frequency (2.76% in controls vs. 2.14% in cases, p = 0.0159, OR = 0.771; 95% CI [0.624-0.949], Fisher’s exact test, Supplementary Table S5), suggesting a potential protective effect of OCT3 coding variants against psychiatric disease. The combined group of coding variants encompasses potential ‘loss of function’ (LoF), non-functional, and potential gain of function variants of varying effect sizes. A separate carrier-based association analysis of the variants that completely disrupt OCT3 function, i.e. the identified nonsense, frameshift and splice site variants revealed an even more pronounced enrichment in control subjects relative to cases, with carrier frequencies of 0.512% and 0.233% respectively (p = 0.0057, OR = 0.454; 95% CI [0.268-0.788], Fisher’s exact test). For the remaining group of coding missense variants, we observed an interesting tendency for accumulation in control individuals (p = 0.149, OR = 0.846; 95% CI [0.846-1.062], Fisher’s exact test, Supplementary Table S5). In light of these genetic data, we performed detailed functional investigations of 24 selected missense variants to interrogate the structure-function relationship of OCT3 (Fig. 4b, Supplementary Table S6).

### Structure-function analysis of OCT3 genetic variants

The herein investigated genetic variants^10,12,35-40^ were mapped on the structure of OCT3 (Fig. 4a) and functionally characterized *in vitro* (Fig. 4b, Supplementary Fig. 14-17, Supplementary Table S6). While some of the OCT3 mutants displayed protein expression or cell surface trafficking defects, many reached at least 60% of wild-type surface expression but showed reduced uptake function (Fig. 4b). A classification of the function-perturbing mutations suggests the existence of three classes: (a) mutations directly affecting the substrate translocation pathway (indicated by reduced uptake but conserved surface expression), (b) mutations modulating the conformational transitions (reducing uptake) (c) mutations affecting protein folding and cell surface expression (indicated by lower surface expression measured by confocal microscopy and higher Förster Resonance Energy Transfer (FRET) due to intracellular accumulation of proteins).

**Figure 4.**
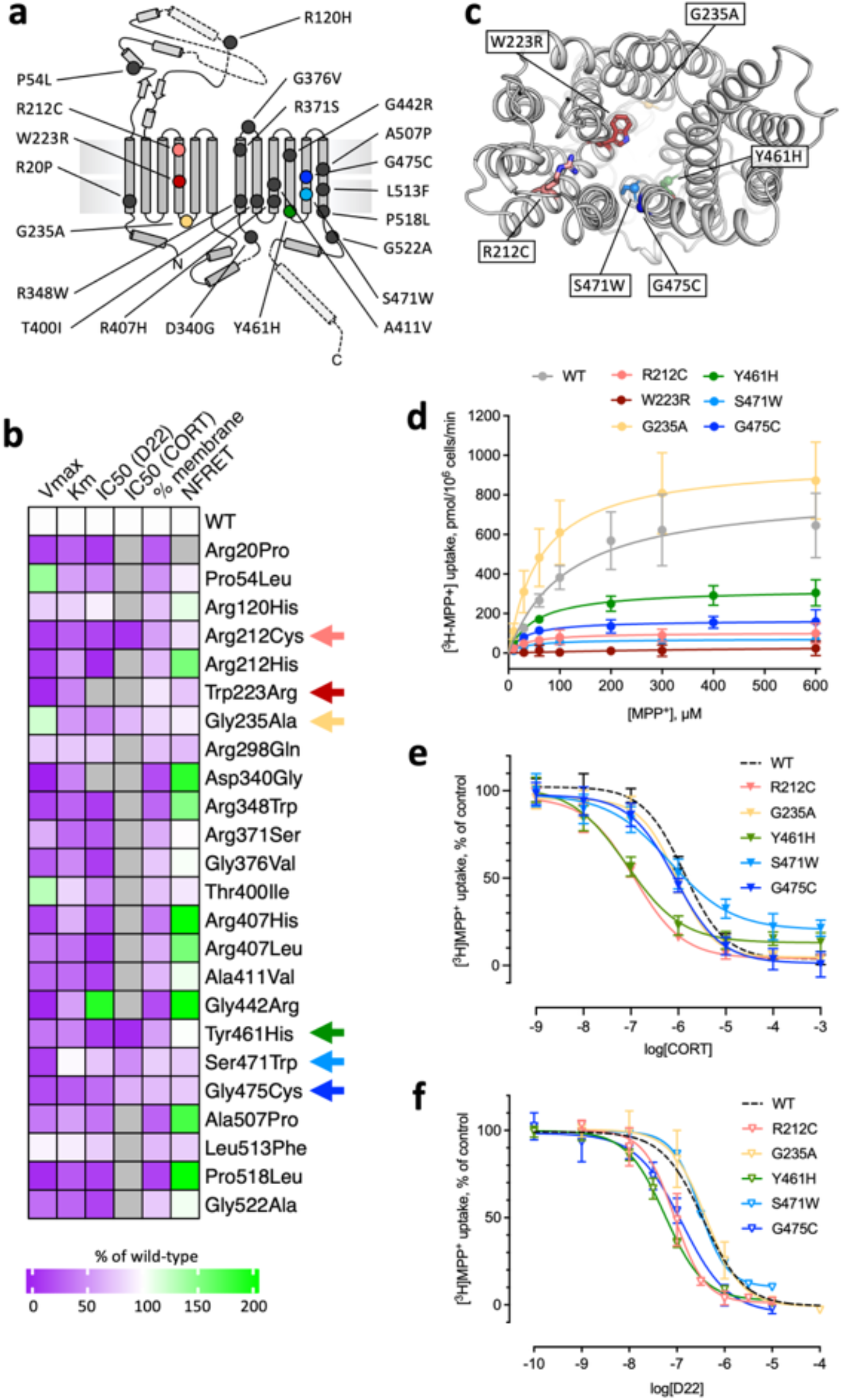
Functional analysis of OCT3 mutants. **a**, Distribution of the OCT3 genetic variants, mapped on the structure of OCT3 **b**, A heat map illustrating the effects of mutations. The scale bar indicates the increase (green) or decrease (magenta) of V_max_, K_m_, IC_50_, membrane expression and NFRET compared to wild-type, as detailed in Materials and Methods. Arrows indicate the variants selected for detailed characterization. **c**, View of OCT3 from the extracellular space, with the side-chains of the selected residues shown as sticks. **d**, Uptake of MPP^+^ by the wild-type OCT3 (WT) and by the selected variants expressed in HEK293 cells (Table S5). **e-f**, Inhibition of MPP^+^ transport by CORT (**e**) and D22 (**f**). The WT is indicated with a dotted line. The values in d-f correspond to mean ± SD, n = 3-4; in triplicate.

The structure of OCT3 allows for a rational interpretation of the large collection of functional data. This can be summarized as follows: (i) Genetic variants located in the ECD of OCT3 do not or barely affect uptake velocity and affinity for ligands; in fact, some variants even have higher V_max_- and K_m_-values than OCT3-WT (e.g. P54L, R120H). (ii) Genetic variants located in proximity of residue 340 interfere with folding and cause ER retention, with only little or no surface expression of D340G and R348W. (iii) Genetic variants located in the ICL (G235A, R298Q, with the exception of D340G) do not affect transport and ligand affinity. (iv) Genetic variants located in TM1, 5, 7, 9, 10, 11 and 12 mostly led to reduced transport velocity; (v) genetic variants near the translocation pathway (TM4, TM11) have pronounced effects on uptake and affinity (Fig. 4a-b, Supplementary Fig. 14-17, Supplementary Table 6).

Subsequently, we focused on mutations directly affecting substrate translocation and selected the six mutants in immediate proximity of the substrate binding site or the translocation pathway (Fig. 4a-c). Functional assays, including MPP^+^ uptake and D22 and CORT uptake inhibition assays, revealed that three of the six selected mutants stand out: R212C, W223R and Y461H (Fig. 4d-f).

Substitution of W223 by arginine did not impair delivery of the protein to the cell surface but resulted in an inactive transporter (Supplementary Fig. 18). Residue W223 is central to the substrate binding site, making extensive interactions with CORT. Substitution of W223 by R removes the large aromatic sidechain and places a positive charge into the substrate binding site. This is likely to interfere with binding of organic cations by electrostatic repulsion. Simulations showed that the environment of W223, exemplified by the distances of W223 to Y227 and Q247, is strongly dependent on the presence of the ligands within the binding site (Fig. 5a-b). Thus, it is likely that W223R mutation does not only affect protein-substrate interactions, but also influences the neighboring residues within the substrate translocation pathway.

**Figure 5.**
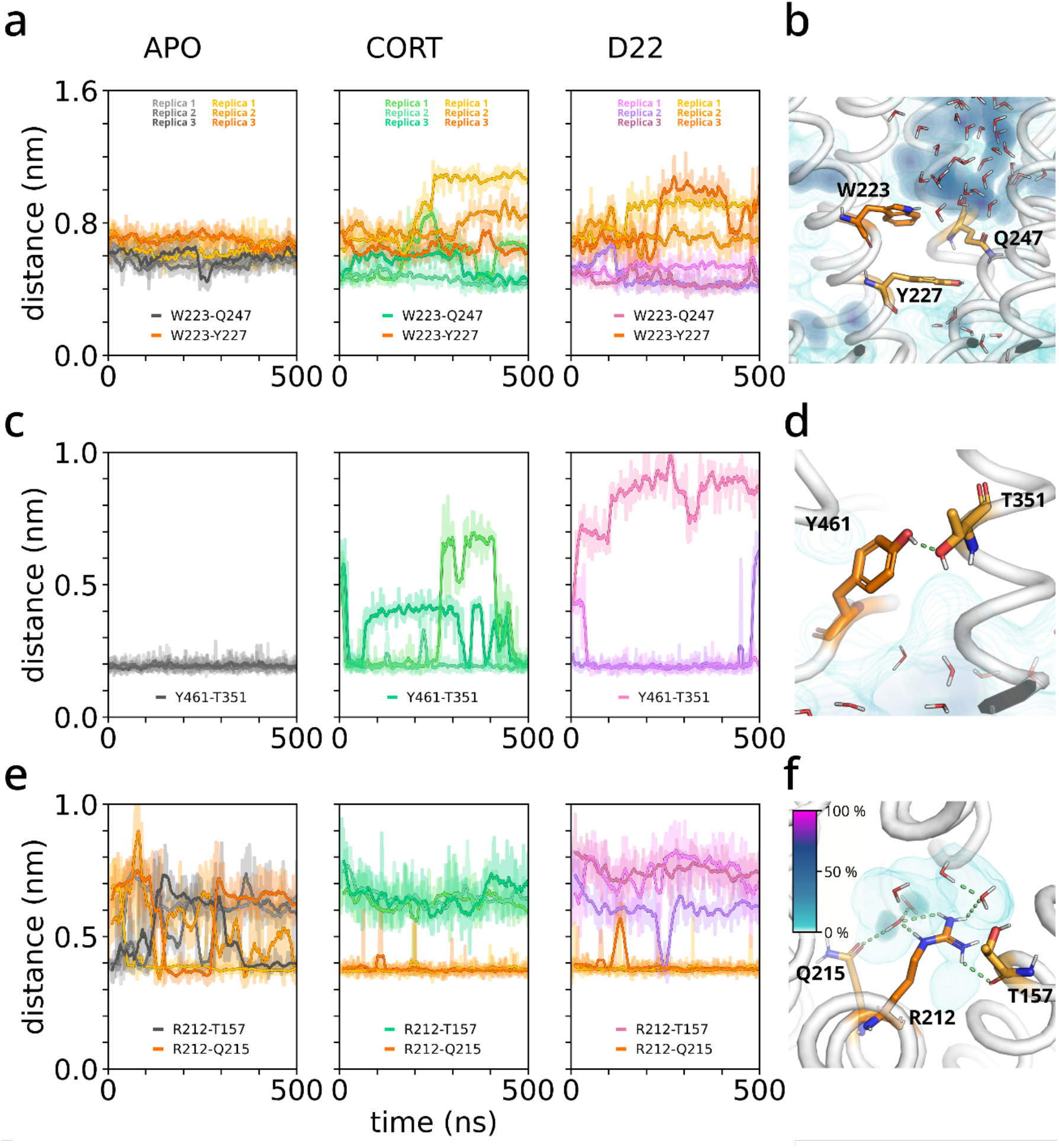
The possible underlying causes of function disruption in genetic variants of OCT3. **a**, Distance between the side chains of W223 and Y227 or Q247, indicating the orientational dynamics of the sidechain of W223. **b**, Local environment of W223, showing average water molecules occupancy in its proximity. **c**, Donor-acceptor hydrogen bond distance between Y461 and T351, confirming the long-range effect of the bound inhibitor on the local structural stability of the Y461 site. **d**, Zoom onto Y461, showing the hydrogen bond to T351 an averaged water occupancy in its proximity. **e**, Distance of the side chain of R212 to residue T157 and G215, highlighting the oscillation of the interaction pattern in the apo state compared to the stable conformation in the inhibitor bound states. **f**, Close-up of R212, emphasizing on the hydrogen bond network formed with residues Q215, T157 and water molecules. Averaged spatial occupancy of water is represented as a density colored according to the legend.

The mutants R212C and Y461H both displayed reduced V_max_ and lower K_m_ for transport of MPP^+^ and showed higher affinity for the inhibitors D22 and CORT (Fig. 4d-f, Supplementary Table 3). These two residues are both located within the hydrophobic core but on opposite sides, i.e. R212 (TM4) in the vicinity of the extracellular vestibule and Y461 (TM10) close to the intracellular end of the substrate permeation pathway (Fig 4a,c) are sandwiched between TM1, TM2, TM3, TM4, and TM6 and between TM7, TM10 and TM11, respectively. Both residues are shielded from the substrate translocation path and may have a role in stabilizing TM4 (R212) and TM10 (Y461), which line the substrate translocation pathway.

Residue Y461 is located in a hydrophobic pocket and forms a very stable hydrogen bond with T351 on TM7 (Fig. 5c-d), thereby fixing the distance between TM7 and TM10. Binding of the inhibitors D22 and CORT appears to destabilize this hydrogen bond, based on MD simulations (Fig. 5c-d). The Y461H mutation is likely to break the hydrogen bond and through its much higher polarity attract water molecules. Together these changes may destabilize the central helix TM10. It is conceivable that Y461 plays a role in maintaining the structural integrity in the whole domain-based conformational transitions of OCT3 during the transport cycle.

The side chain of R212 is in proximity of the main-chain atoms of L35, V39 (TM1), T157 (TM2) and the side chains of T157 (TM2), Q215 (TM4), and Q251 (TM6) (Fig. 5e-f). TM1, TM2 and TM4 contain residues involved in substrate/inhibitor interactions: mutation of R212 to a cysteine led to a reduction in transport V_max_ and a decrease in K_m_. This is consistent with the assumption that interactions with R212 are required for an effective transport cycle. Our MD simulations show that R212 participates in an extended hydrogen bonding network, which includes the nearby polar residues and 2-4 water molecules that fill the intra-membrane cavity surrounding R212 (Fig. 5e-f). In apo OCT3, this network is very dynamic, as R212 cannot establish all interactions simultaneously. By slightly shifting its position, R212 interacts directly with the side chains of T157, Q215 and Q271, while the additional interactions are formed through a water bridge. Binding of the inhibitors D22 or CORT induces small structural changes in OCT3, reducing the conformational dynamics of R212 as it partitions towards Q215 and Q271. The differential effects of the ligands on R212 environment and dynamics, combined with the functional studies of the R212 mutants, suggest an important role of this residue in controlling the conformational changes of OCT3 during ligand binding and translocation.

## Discussion

The SLC22 family comprises more than 30 transporters, which facilitate the transport of organic cations (OCTs), anions (OATs) and zwitterions (OCTNs)^1^. Collectively, these transporters define the pharmacokinetics of a vast array of drugs and xenobiotics^41,42^. Herein, we describe the cryo-EM structure of OCT3 and provide the first direct insights into the organization of a SLC22 member, its substrate permeation pathway and ligand binding pocket. Both ligands of which we herein report cryo-EM structures, are handled by OCT3 in different ways which only partially overlap. It is not surprising, however, that the binding site of OCT3 allows accommodation of many diverse binding partners; the behavior rather substantiates the poly-specificity of a class of transporters which interact with a wide and complex array of compounds: from the antiviral drug abacavir and the antidiabetic drug metformin to the antineoplastic drug sunitinib^1^.

However, the current consensus in the field is that both corticosterone and D22 are non-transported inhibitors with scarce evidence that D22 may accumulate into astrocytes via an OCT3 dependent mechanism^43^. It remains to be determined whether OCTs are able to move inhibitors across the membrane. This property would be similar to multidrug transporters, such as ABCB1 and ABCG2, which are capable of transporting a variety of organic compounds, including their inhibitors elacridar and tariquidar^44^. Our structures show that OCT3 inhibitors may sterically block the translocation pathway. Future studies will be necessary to determine whether these poses of the inhibitors completely impede transport, or whether the transporter is nevertheless capable of moving the inhibitor molecules across the lipid bilayer with very low efficiency.

Several questions remain, in particular concerning the substrate translocation pathway of OCT3 and the role of different domains in its activity. OCT3 shows an obvious access path to the substrate binding site that is wide open to the extracellular side. However, some of the substrates of OCT3 are hydrophobic, which may suggest accumulation in the lipid bilayer. It is tempting to speculate that OCT3 might be able to retrieve its substrates from the membrane, which is a mechanism well described for ABC transporters^45^. The OCT3 structure provides a hint for the location of such a lateral access site (the V-site), which remains to be validated. The role of the ECD of OCT3 is currently unexplored, and the mutations located in the ECD did not substantially affect transport properties of OCT3. However, by analogy with other MFS transporters that contain an extended ECD^46,47^ this domain in OCTs may play an important role in molecular gating or in protein-protein interactions. For example, ECD could play a role in recruitment of CD63, a binding protein known to interact with OCTs, and to regulate their trafficking to the plasma membrane or intracellular compartments^48^. Moreover, it has been shown that both OCT1 and OCT2 depend on an intact ECD for oligomerization and trafficking to the plasma membrane^49,50^.

The extended list of novel genetic variants functionally characterized allows us to place missense mutations with clear functional consequence in the context of an experimentally determined OCT3 structure. Genetic variants located in certain regions of the protein clearly impact transport and/or surface expression of OCT3 and thus impact uptake of physiological substrates and drugs. Moreover, as new therapeutic strategies involving OCTs emerge in a number of different areas, a better understanding of how genetic variations affect transporter function may be useful to understand personalized variations in drug response and to develop effective pharmacogenomics-based therapeutic strategies. Thus, the deeper knowledge of the structure and function of OCTs brings us closer to more precise and efficient medical treatments^22,41,42^. Therefore, our structures provide a new starting point for rational drug development targeting OCT3 in the treatment of depression^22,51^, diabetes^11,12,52^, cardiac disease^10,15^, and cancer chemotherapy^19,39,53^.

## Methods

All methodological details can be found in the supplementary material.

## Data availability

Data supporting the findings of this study are available within the article and its Supplementary Information Files and from the corresponding authors upon reasonable request.

## Acknowledgements

The research described in this publication was supported by the Vienna Science and Technology Fund (WWTF) [CS 15–033] (HHS), Austrian Science Fund (FWF) doctoral program Molecular Drug Targets [W1232] (HHS), stand-alone project P34670-B20 (HHS, TS and VMK) and doctoral program Neuroscience [DOC33-B27] (HHS and JM), Theodor Körner Fonds 2020 (JM), Swiss National Science Foundation (SNSF; grant No. P400PM_191032, DL; Sinergia grant No. 198545, VMK), and the Lundbeck Foundation (R303-2018-3540, FH). This project has received funding from the European Union’s Horizon 2020 research and innovation programme under the Marie Skłodowska-Curie grant agreement No 860954 (TS).

We thank Michael Freissmuth and Richard Kammerer for critical comments on the manuscript. In addition, we thank the PSI EM Facility for their support: Emiliya Poghosyan and Elisabeth Müller-Gubler, as well as Miroslav Peterek and Bilal Qureshi (ScopeM, ETH Zurich) for their support in cryo-EM data collection. We also thank Pavel Afanasyev (CEMK, ETH Zurich) for his advice and help in cryo-EM data handling. In addition, we thank Spencer Bliven and Marc Caubet Serrabou (PSI) for their support in high performance computing. In addition, the results presented have, in part, been achieved using the Vienna Scientific Cluster.

The authors wish to dedicate the present work to Heinz Bönisch.

## Author contributions

B.K., J.M., F.H., R.G., E.L., D.L., T.S., V.M.K., H.H.S. designed experiments, analysed data and prepared figures. B.K., J.M., F.H., R.G., E.L., D.L. J-W.Y., M.H., K.J., O.K., K.S. performed experiments and analysed data. T.W. contributed to conception and critical revisions. U.G., T.S., V.M.K. and H.H.S. supervised the project. B.K., J.M., T.S., V.M.K. and H.H.S. wrote the manuscript with input from all co-authors.

## Competing interests

The authors declare no competing interests.

## Notes

### Competing Interest Statement

The authors have declared no competing interest.

### Summary of Updates

The new version contains new data (time dependency, new uptake inhibition data at OCT1), extended simulation time (now 1 micro second) and improved figures of the structural part.

