## Supplementary material for "Structural basis of organic cation transporter-3 inhibition"

### MATERIALS AND METHODS

#### Materials and chemicals

Detergents, dodecylmaltoside (DDM) and glyco-diosgenin (GDN), and Brain Polar Lipids were purchased from Antrace Inc. Decynium-22 was ordered from Synthon Chemicals (Bitterfeld-Wolfen, Deutschland). All other chemicals and cell culture supplies were obtained from Sigma-Aldrich (St. Louis, MO, USA) and Sarstedt (Nuembrecht, Germany), unless indicated otherwise.

#### Cell lines and cell culture

hOCT3 wildtype plasmid was generously provided by Eric Gouaux, Vollum institute, Oregon Health & Science University, Portland, OR, USA. Fluorescently tagged constructs (YFP and CFP) were generated by N-terminal fusion of the fluorescent protein to the plasmid in a pcDNA3 vector. QuikChange site-directed mutagenesis kits and QuikChange Primers (Agilent Technologies, Santa Clara, USA) were used to create plasmids of hOCT3 single nucleotide polymorphisms. Constructs were verified by sequencing (LGC Genomics, Berlin, Germany). Stable polyclonal HEK293 cell lines, expressing hOCT3 wildtype or hOCT3 genetic variants, were established as follows: HEK293 cells were transfected with the respective plasmids with the jetPRIME transfection method, according to the manufacturer's instructions (VWR International GmbH, Vienna, Austria). High selection pressure was maintained for ten days by adding 100 µL geneticin (G418, 50 mg x $mL^{-1}$). 500,000 cells were then FACS sorted and polyclonal cell lines established, according to expression levels.

In cell culture, cells were maintained in high glucose- (4.5 g x $L^{-1}$), l-glutamine-containing (584 mg x $L^{-1}$) Dulbecco's Modified Eagle Medium (Sigma-Aldrich, St. Louis, USA) with 10 % heat-inactivated Fetal Bovine Serum (FBS, Sigma-Aldrich), penicillin (1 U x $mL^{-1}$, Sigma-Aldrich) and streptomycin (1 µg x $mL^{-1}$, Sigma-Aldrich) added. For maintaining the selection process, geneticin (50 µg x $mL^{-1}$) was added regularly. Cells were maintained in 10-cm cell culture dishes (Greiner) at 37 °C and 5 % $CO_2$ in an incubator. The day before uptake and uptake inhibition assays, cells were seeded onto PDL (poly-D-lysine) coated 96-well plates at a density of approximately $0.4 \times 10^5$ cells per well. For live confocal microscopy, cells were seeded onto PDL-coated 35mm glass-bottom dishes (Cellvis, Sunnyvale, California, USA) at a density of $0.3 \times 10^6$ cells per dish.

#### Radiotracer uptake assays

On the day of the experiment, cells were incubated with 50 µL Krebs-HEPES-buffer (KHB; 10 mM HEPES, 120 mM NaCl, 3 mM KCl, 2 mM $CaCl_2$, 2 mM $MgSO_4$ and 20 mM D-glucose, pH adjusted to 7.3) containing increasing concentrations of 1-methyl-4-phenylpyridinium ($MPP^+$; Sigma-Aldrich, St. Louis, MO, USA) together with 50 nM $[^3H]$-$MPP^+$ (80–85 µCi $mmol^{-1}$; American Radiolabeled Chemicals, St. Louis, USA) for ten minutes. Time-dependent uptake was determined using 50 nM $[^3H]$-$MPP^+$ for the time indicated. Unspecific uptake was determined in the presence of 100 µM decynium-22. Cells were washed with KHB and then lysed with 100 µL 1 % sodium dodecyl sulfate (SDS). Lysate was transferred to a counting vial, containing 2 mL of scintillation cocktail. Uptake of tritiated substrate was assessed with a beta scintillation counter (Perkin Elmer, Waltham, USA).

#### Radiotracer uptake inhibition assays

Cells were preincubated with vehicle or increasing concentrations of decynium-22 or corticosterone (dissolved in DMSO), diluted in 50 µL of KHB for ten minutes. Preincubation solution was aspirated and uptake solution, containing KHB, vehicle or substance of interest at the desired concentration and 50 nM [$^3$H]-MPP$^+$, was added to the wells. Uptake was terminated by aspiration and cell washing with 100 µL ice-cold KHB after ten minutes. Cells were subsequently lysed with 100 µL of 1 % SDS. Lysate was transferred to a counting vial, containing 2 mL of scintillation cocktail. Uptake of tritiated substrate was assessed with a beta scintillation counter (Perkin Elmer, Waltham, USA).

##### Confocal microscopy and image analysis

Confocal microscopy images were taken on a Nikon A1R+ laser scanning confocal microscope system with a 60× NA 1.4 oil immersion objective (Nikon, Vienna, Austria). Cell culture medium was aspirated and cells incubated with trypan blue (0.4%, Sigma Aldrich) for 10 minutes and then washed with KHB multiple times. Cells were kept on KHB throughout the experiment.

eYFP Fluorescence was excited with a 488 nm, trypan blue with a 561 nm laser line. Emitted light was filtered, using a 525/50 nm (eYFP) or 595/50 nm (trypan blue) emission filter respectively and detected by a high-sensitivity GaAsP PMT detector. Three images were taken on three separate days. Image analysis was conducted in Fiji ImageJ 1.53c[1]. For analysis of membrane expression, in each image two regions of interest were drawn per cell, one encompassing the cell membrane (as defined by trypan blue staining) and one the cell interior. Membrane expression levels were defined as the relative values of membrane versus intracellular mean intensity.

##### Immunoblotting

Cell lysates from heterologous HEK293 cells stably expressing YFP-tagged OCT3 proteins were solubilized in lysis buffer containing 1% Triton X-100, 20 mM Tris-HCl pH 8.0, 150 mM NaCl, 1 mM EDTA, 1 mM sodium orthovanadate, 5 mM sodium fluoride, 5 mM sodium pyrophosphate, and a protease inhibitor mixture (Roche Applied Science) on a tube rotator for 2 hour at 4 °C. After centrifugation at 14,000 x g for 30 min at 4 °C, the supernatant was collected for immunoblotting. Proteins were separated on SDS-PAGE gels and transferred onto nitrocellulose membranes (GE healthcare), which were then blocked and immunostained with a rabbit anti-GFP polyclonal antibody (A6455, Thermo Fisher).

##### Förster Resonance Energy Transfer imaging

Förster Resonance Energy Transfer (FRET) microscopy was utilized to assess protein-protein-interaction of OCT3 isoforms. Experiments were conducted using HEK293 cells transiently expressing OCT3-constructs labeled with a fluorescence donor (eCFP) and acceptor (eYFP) attached to the N-terminus, respectively. The cells were seeded into 29 mm dishes with 20 mm bottom (# 1.5 glass; Cellvis, Mountain View, CA, USA) at a density of $10^5$ cells per dish one day prior to imaging. FRET was measured with an iMIC inverted microscope (T.I.L.L. Photonics GmbH, Kaufbeuren, Germany) equipped with a 60X (N.A. 1.49) oil objective (Olympus).
Fluorescence was excited with a 100 W Xenon Lamp (Polychrome, T.I.L.L Photonics GmbH, Kaufbeuren, Germany). The excitation light was filtered through 436/20 nm (eCFP) or 514/10 nm (eYFP) excitation filters (Semrock, Rochester, NY, USA) and directed to the sample by a 442/514 dual line dichroic mirror (Semrock, Rochester, NY, USA). The emitted fluorescence light was filtered through a 480/40 nm and 570/80 nm dual emission filter (Semrock, Rochester, NY, USA) to a beamsplitter unit (Dichrotom, T.I.L.L Photonics, Kaufbeuren, Germany). Emission light was separated according to fluorescence wavelength using a 515 nm dichroic mirror and channels (<515 nm & >515 nm) projected side by side onto an EMCCD chip (iXon Ultra 897 Andor, Andor Technology, Belfast, UK). Live Acquisition software (version

2.5.0.21; T.I.L.L Photonics GmbH, Kaufbeuren, Germany) was used for recording. Two images were taken per set (donor and acceptor emission after donor excitation and acceptor emission after acceptor excitation). Per condition, ten sets were recorded on each of three experimental days and images were then analyzed using Offline Analysis software (version 2.5.0.2; T.I.L.L Photonics GmbH, Kaufbeuren, Germany). Background fluorescence was subtracted from each image, and one region of interest (part of the plasma membrane) per cell was selected in the CFP channel. The average intensity of each region of interest was used for calculations. HEK293 cells expressing a CFP or YFP signal only were used to determine spectral bleed through (BT) for donor (0.57) and acceptor (0.04). Normalized FRET (NFRET) was calculated as follows

$$\mathrm{NFRET} = \frac{I_{FRET} - BT_{Donor} \times I_{Donor} - BT_{Acceptor} \times I_{Acceptor}}{\sqrt{I_{Donor} \times I_{Acceptor}}}$$

**Genetic data**

Exome sequencing data were provided by the integrated psychiatric research (iPSYCH) consortium. 1,472,762 Danish individuals born between May 1, 1981, and December 31, 2005 were genotyped, which was previously described in detail[1]. The exome sequencing data used in this study is from the integrated psychiatric research (iPSYCH) consortium's first phase genotyping of a nation-wide Danish birth cohort which has been described in greater detail previously[2]. The iPSYCH study sample is approved by the Danish Data Protection Agency. Informed consent is not required by law for register-based research in Denmark. Procedures for exome sequencing, sample and variant quality control was performed as described in[3].

**Data and statistical analysis**

$IC_{50}$, $V_{max}$ and $K_m$ values were calculated and graphs plotted with GraphPad Prism 9.2.0 (GraphPad Software Inc., San Diego, USA). Half maximal inhibitory concentrations ($IC_{50}$) were determined by non-linear regression, solving equation Y=Bottom + (Top-Bottom)/(1 + 10^(X-LogEC50)). Michaelis-Menten kinetics were determined via Y=Vmax*X/(Km+X). All data are from at least three biologically independent experiments (n >= 3), in triplicate and portrayed as mean ± SD. Confocal and FRET microscopy images were taken and Western blot lysates prepared from three individual passages on three separate days. The heatmap was created in RStudio (2021.09.2), package: ComplexHeatmap. Fisher's-exact test was used to compare carrier frequencies of coding SLC22A3 variants in cases and controls.

**Protein expression and purification**

The full-length human OCT3 (Uniprot: O75751) was cloned into pACMV-based vector with C-terminal 3C-YFP-TwinStrep fusion tag. The plasmids were transfected into HEK 293F cells and a stable monoclonal cell line (HEK293F-hOCT3) capable of expressing OCT3 was generated for large scale expression. HEK293F-hOCT3 cells were grown in suspension at 37 °C to a density of ~3-3.5x $10^6$ $mL^{-1}$ in Gibco® FreeStyle™ 293 Expression Medium (ThermoFisher Scientific). The cells were collected by centrifugation at 800 g for 20 minutes and stored at -80 °C until the day of the experiment.

For purification, the cell pellets collected from to 0.5 L of cell culture were thawed and resuspended in buffer A (50 mM Tris-HCl, 150 mM NaCl, pH 8.0) supplemented with protease inhibitors (1 mM benzamidine, 1 µg/mL leupeptin, 1 µg/mL aprotinin, 1 µg/mL pepstatin, 1 µg/mL trypsin inhibitor and 1 mM PMSF) and lysed using a Dounce homogenizer. The lysate was centrifuged at 186000 x g (Ti45 rotor) for 45 min, and membrane pellet was solubilized in the buffer B (50 mM Tris-HCl, 150 mM NaCl, 10 % glycerol, 1 % dodecylmaltoside (DDM) and 0.02 % cholesteryl hemisuccinate (CHS), pH 8.0) for 1 hour at 4 °C. The lysate was cleared by ultracentrifugation (Ti45 rotor, 186000 x g for 30 min to remove insoluble debris. The supernatant was incubated with 4 mL of CNBr-activated Sepharose coupled to an anti-GFP nanobody[4]. After a 60 min incubation at 4 °C the resin was collected in a gravity column and washed with 40 column volumes of buffer C (50 mM Tris-HCl 150 mM NaCl, 0.02 % glycol-diosgenin (GDN), 5 % glycerol, pH 8.0). The protein was eluted using cleavage by HRV 3C protease (1:10 w/w). The eluted protein was concentrated with a 30 kDa cut-off Amicon Ultra spin (Millipore) and subjected to size-exclusion chromatography (SEC) using a Superpose 6 Increase 10/300 GL column (GE Healthcare) equilibrated in buffer D (50 mM Tris-HCl, 150 mM NaCl, 0.02 % GDN, pH 8.0). The fractions corresponding to purified hOCT3 (elution volume, 14.8-16.9 mL) were collected and pooled. The protein purity was assessed using SDS-PAGE.

GFP-nanobody: The expression and purification of GFP nanobody was carried out as previously described[4]. In short, the anti-GFP nanobody plasmid was transformed into *E. coli* BL21 (DE3) and grown in Luria Broth (LB) media supplemented with 50 µg/ml ampicillin at 37 °C. Once the optical density at 600 nm (OD600) of bacterial culture reached 0.5, the expression of protein was induced by adding 0.5 mM IPTG followed by overnight incubation at the 20 °C. Bacterial cultures were harvested at 4,000 g for 20 min at 4 °C, and pellets were frozen in liquid N2 and stored at −80 °C. Frozen pellets were thawed on ice, re-suspended in buffer E (25 mM HEPES pH 8.0, 150 mM NaCl, 10 mM Imidazole, 1 mM PMSF, 10 ug/mL DNase I). The cells were lysed by sonication, and the cell lysate was centrifuged for 30 min at 20000xg. Clarified lysate was then incubated with Ni-NTA resin (1-2 mL bed volume of resin per 1 L of culture) for 30-40 minutes, and subsequently washed with 20 CV of buffer F (25 mM HEPES pH 8.0, 150 mM NaCl) supplemented with 50 mM imidazole, and later protein was eluted with 5 CV buffer F containing 250 mM Imidazole. The elution was concentrated with a 10 kDa cut-off Amicon concentrator and loaded on a Superdex 75 16/600 GL column (GE Healthcare) in buffer F. SEC fraction corresponding to GFP-nanobody was pooled and flash frozen in liquid N2, and stored at −80 °C. Protein concentration was determined by absorbance at 280 nm using ε = 27055 $cm^{-1}M^{-1}$.

Membrane Scaffold Protein: The expression and purification of Membrane Scaffold Protein MSP1D1 was carried out as previously described[5]. In brief, the MSP1D1 plasmid was transformed into *E. coli* BL21 (DE3) and grown at 37 °C in Terrific Broth (TB) media. The expression of protein was induced with 1 mM IPTG at OD600 of ~2-3 and incubated for 3 hour. After harvesting by centrifugation, cell pellets was re-suspended in lysis buffer (50 mM Tris-HCl, 200 mM NaCl, 25 mM Imidazole, 1 % Triton-X100, 1mM PMSF and 10 ug/ml DNase I) and lysed by sonication. The clarified lysate after centrifugation (20000xg for 30 minutes) was incubated with Ni-NTA resin for 30 minutes, and subsequently step washed with 10 CV of buffer G (50 mM Tris-HCl pH 8.0, 150 mM NaCl) containing 25 mM Imidazole, 1 % Triton-X100, 5 CV of buffer H (50 mM Tris-HCl pH 8.0, 150 mM NaCl, 25 mM Imidazole, 2 % Sodium Cholate), 5 CV of buffer I (50 mM Tris-HCl pH 8.0, 150 mM NaCl, 50 mM Imidazole). Finally the protein was eluted in buffer I supplemented with 350 mM Imidazole. Elution fractions containing MSP1D1 were pooled, desalted in buffer J (20 mm Tris-HCl pH 8.0, 200 mM NaCl) and flash frozen in liquid N2, and stored at -80 °C. Protein concentration was determined by absorbance at 280 nm using ε = 21430 $cm^{-1}M^{-1}$.

**Reconstitution in lipid nanodisc**

Reconstitution of OCT3 in MSP1D1 nanodiscs was performed using freshly purified transporter. In brief, 2.5 mg of brain polar lipid (BPL, Avanti) dissolved in 100 µL of chloroform was dried under a stream of $N_2$. Dried film of lipid was mixed with 300 µL of 3 % DDM and the lipid-detergent mixture was then sonicated using bath sonicator (Bandelin SONOREX™ SUPER, Germany) until the mixture turned translucent. Detergent-solubilized BPL extract was added to freshly purified OCT3 at a molar ratio of 1 : 75, and incubated for 30 min at room temperature with rotation. MSP1D1 was added to protein-lipid mixture and incubated for additional 30 min. Concentrations of OCT3 for nanodisc reconstitution were in the 10-15 μM range, with a molar ratio of OCT3 to MSP1D1 to lipid of 1:1:75. Following the incubation period, nanodisc formation was triggered by adding 300 mg of wet Bio-beads (washed with 100 % methanol and with Milli-Q water). This final reconstitution mixture was incubated at 4 °C for 16 h (overnight) with gentle mixing. The supernatant was cleared of the beads by letting the beads settle and removing liquid carefully with a pipette. Sample was spun for 10 min at 25000 x g using a bench-top Eppendorf centrifuge before loading onto a Superose 6 Increase 10/300 GL column equilibrated in 20 mM Tris-HCl, pH 8.0, 150 mM NaCl. The peak fractions corresponding to OCT3 in MSP1D1 (elution volume, 15.4-16.9 mL) were collected, concentrated with a 30 kDa cutoff Amicon concentrator and used for cryo-EM grid preparation.

##### Cryo-EM sample preparation and data collection

The cryo-EM samples were prepared using freshly reconstituted OCT3-BPL-MSP1D1 nanodisc samples. The complexes of OCT3-D22 and OCT3-Corticosterone were prepared by adding D22 (100 mM stock concentration in 100 % DMSO) or CORT (50 mM stock concentration in 100 % DMSO) at a final concentration of 1 mM, and incubating the samples on ice for 10-15 minutes. A final concentration of OCT3 and OCT3-drug complexes used for freezing grids was ~6-7 mg/mL, estimated based on absorbance at 280 nm and normalized extinction coefficient of 133700 $M^{-1}$ $cm^{-1}$ (considering two molecules of MSP1D1 per one OCT3 molecule).

Quantifoil 1.2/1.3 grids (300 mesh) were briefly glow-discharged in a PELCO easiGlow (Ted Pella) glow discharge cleaning system, for 25 sec at 30 mAmp in air. A small amount of protein or protein-drug sample (3.5 μL) was then applied to glow-discharged grids, blotted for 3 sec with blot force 20, and plunged into liquid ethane using Vitrobot Mark IV (Thermo Fisher Scientific) with 100 % humidity at 4 °C. The frozen grids were transferred under cryogenic conditions and stored in liquid nitrogen for subsequent cryo-EM data collection.

##### Cryo-EM data collection and processing

A total of four datasets with 5580, 12251, 6191, 5227 movies were collected using EPU on a 300 kV Titan Krios (Thermo Fisher Scientific) equipped with a Gatan K3 direct electron detector and a Gatan Quantum-LS GIF at ScopeM, ETH Zurich. All movies were acquired in super-resolution mode with a defocus range of -0.5 to -3 μm and a final calibrated pixel size of 0.33 Å. The total dose per movie was 48, 48, 48 and 49 e-/Å$^2$ for datasets 1, 2, 3, and 4, respectively. The cryo-EM processing was performed in Relion 3.1.2[6]. The flow chart of cryo-EM processing of apo-OCT3 is depicted in Figure S1. In brief, all movie stacks were motion-corrected using MotionCorr 1.1.0[7] and binned two-fold. All micrographs were CTF corrected using Gctf[8]. A total of 513 particles were manually picked and 2D classified to generate autopicking templates. Selected 2D classes from manually picked particles were used to autopick particles from all micrographs in dataset 1. After multiple rounds of 2D clarifications, the best set of 2D classes were used to generate an initial model to be used in 3D classification. The 3D projections from the best 3D class from dataset 1 were used as templates for autopicking of particles in the datasets 2, 3 and 4. After multiple rounds of 2D and 3D clarifications, a set of 316323 particles was selected for masked 3D refinement (using masks including or excluding the nanodisc density), resulting in a 3D reconstruction at 3.65 Å resolution. The refined particles were subjected to another round of 3D classification without alignment and with masking of the nanodisc. The particles from the best 3D class were subjected to several iterative cycles of 3D refinement, CTF refinement and particle polishing, yielding a final post-processed density map at 3.2 Å resolution.

The data collection strategy and image processing of OCT3-drug complexes was similar to that of OCT3 apo. For the OCT3-D22 complex, three datasets of 11340, 15456, 29004 movies were collected and processed. The movies for dataset 1 were recorded with a total dose of 51 e-/Å$^2$, dataset 2 and 3 with a dose of 49 e-/Å$^2$. For OCT3-CORT complex, a total of three datasets with 23504, 10988, 15035 movies were collected, with total dose of 49, 55, 61 e-/Å$^2$, respectively. The movie stacks of OCT3-D22 dataset 3 and all datasets of OCT3-CORT were binned 2-fold during data acquisition in EPU. The detailed steps of OCT3-D22 and OCT3-CORT processing are shown in Supplementary Fig. 3 and 4, respectively.

##### Model building and refinement

Model building of OCT3 was performed in Coot[9] using the final postprocessed density maps. All maps showed good quality in the membrane-embedded portion of the transporter which, along with a SwissModel[10] generated homology model of the OCT3 TM region (using PDB ID: 6h7d as a template), greatly facilitated model building based on the protein sequence. The ectodomain of OCT3 was partially resolved and thus could not be completely modelled experimentally. A complete hybrid model of OCT3, including the ectodomain and all regions missing in the density map, was built using a model generated by AlphaFold (AF-O75751-F1)[11], joining the missing loops with the model built into the density map. The pLDDT scores for regions modelled using AlphaFold are tabulated in Supplementary Table 2. The ligands, D22 and CORT, were generated from SMILE codes using eLBOW[12]. The structures were refined using phenix.real_space_refine[13]. The quality of the final models were assessed using MolProbity[14]. All figures were generated using PyMOL 2.5.2[15] and ChimeraX[16].

The homology models of OCT1, OCT2 and OAT1 were generated using SwissModel[10] with OCT3 as template. The sequence identity and modelling scores are tabulated in Supplementary Table 3.

##### Molecular dynamics simulations

###### System setup and preparation of the environment

The experimentally solved cryo-EM structure of apo OCT3, as well as the hybrid models that include the AlphaFold generated loops of apo, D22-bound and CORT-bound OCT3 were embedded in a cholesterol-phospholipid containing bilayer. In addition, because of the uncertainty of placing CORT into the cryo-EM density, also models for the second possible orientation of CORT-bound OCT3 structures (CORT-flipped) were created. All glutamate and aspartate sidechains were protonated using their default protonation state, histidine residues were neutralized and protonated at their epsilon nitrogen. To ensure complete lipid mixing and an equilibration of the lipid environment surrounding OCT3, the systems were first simulated using the coarse grained (CG) representation of the MARTINI force field[17]. The simulation box (10.0 x 10.0 x 14.6 nm) harbors water, 150 mM NaCl, 70:30 mol% POPC:cholesterol lipid mixture. OCT3 was embedded using the insane procedure[18] and simulated for 4 µs, while applying position restraints on OCT3. The equilibrated environment was backmapped to an all atom representation[19], while the CG OCT3 was replaced with the original all-atom structure to remove spurious structural distortions introduced by the conversion procedures. The assembled system was relaxed using the membed[20] procedure to relax possible local atom overlaps between OCT3 and its environment. The chain ends at the gaps of the experimentally determined OCT3 structure were neutralized using capping groups (acetylation and methylation) to avoid termini-charge dependent effects.

###### Molecular dynamics simulation

The amber99sb-ildn force field was used to describe the proteins, water and ions[21]. Force field parameters of corticosterone as well as D22 were implemented by applying the general amber force field (GAFF)[22] and ACPYPE[23], partial charges have been calculated by the R.E.D. Server[24]. The membrane was described using slipid[25]. All atom MD simulations were performed using Gromacs 2021.3[26]. All systems were energy minimized and equilibrated in 4 steps that consist of 2.5 ns long simulations, while slowly releasing the position restrain forces acting on the Cα atoms as well as the ligand, if present (1000, 100, 10, 1 kJ/mol/nm). Initial random velocities were assigned independently to every system. Production simulations were performed for 1.0 µs for the apo transporter and 0.5 µs for the ligand bound OCT3, while simulations of CORT-flipped OCT3 were 150 ns long. Temperature was maintained at 310 K using the v-rescale (τ = 0.5 ps) thermostat[27] by separately coupling solvent, membrane, and protein plus ligand, if present. Semi-isotropic pressure coupling was applied using the Parrinello-Rahman barostat[28], using 1 bar and applying a coupling constant of 20.1 ps. Long range electrostatic interactions were described using the smooth particle mesh Ewald method[29] with a cutoff of 0.9 nm. The van der Waals interactions were described using the Lennard Jones potentials applying a cutoff of 0.9 nm. Long range correction for energy and pressure were applied. Coordinates of all atoms were recorded at every 25 ps. The Molecular Dynamics Parameters (MDP) files used during production can be found in the SI.

#### Supplementary Figures

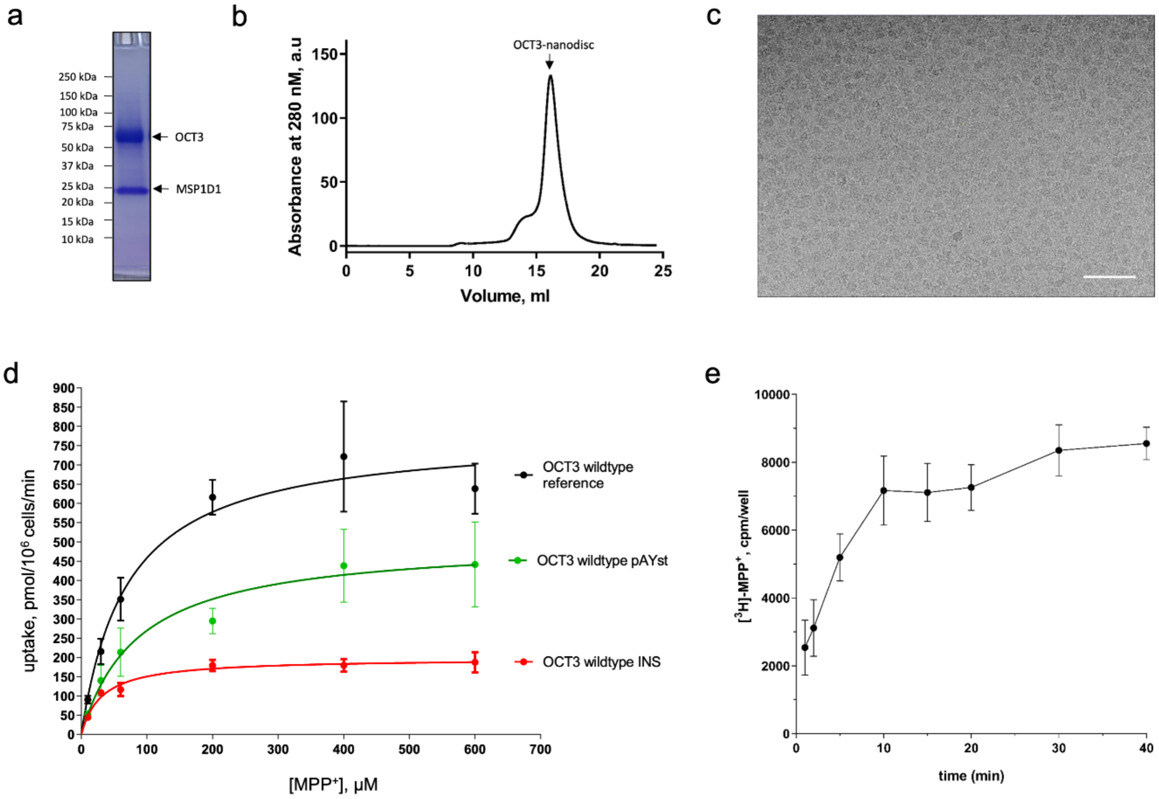


**Supplementary Fig. 1. Expression, purification, and reconstitution of OCT3. a,** SDS-PAGE analysis and **b,** SEC profile of OCT3 reconstituted in MSP1D1. **c,** A representative micrograph (scale bar, 25 nm) of the OCT3-nanodisc sample. **d,** [$^3$H]-MPP$^+$ uptake comparison of HEK293 cells expressing hOCT3 in different vectors (n = 3, in triplicate; error bars denote S.D.). **e,** Time course of [$^3$H]-MPP$^+$ uptake in HEK293 cells expressing YFP-hOCT3 (n = 4, in triplicate; error bars denote S.D.).

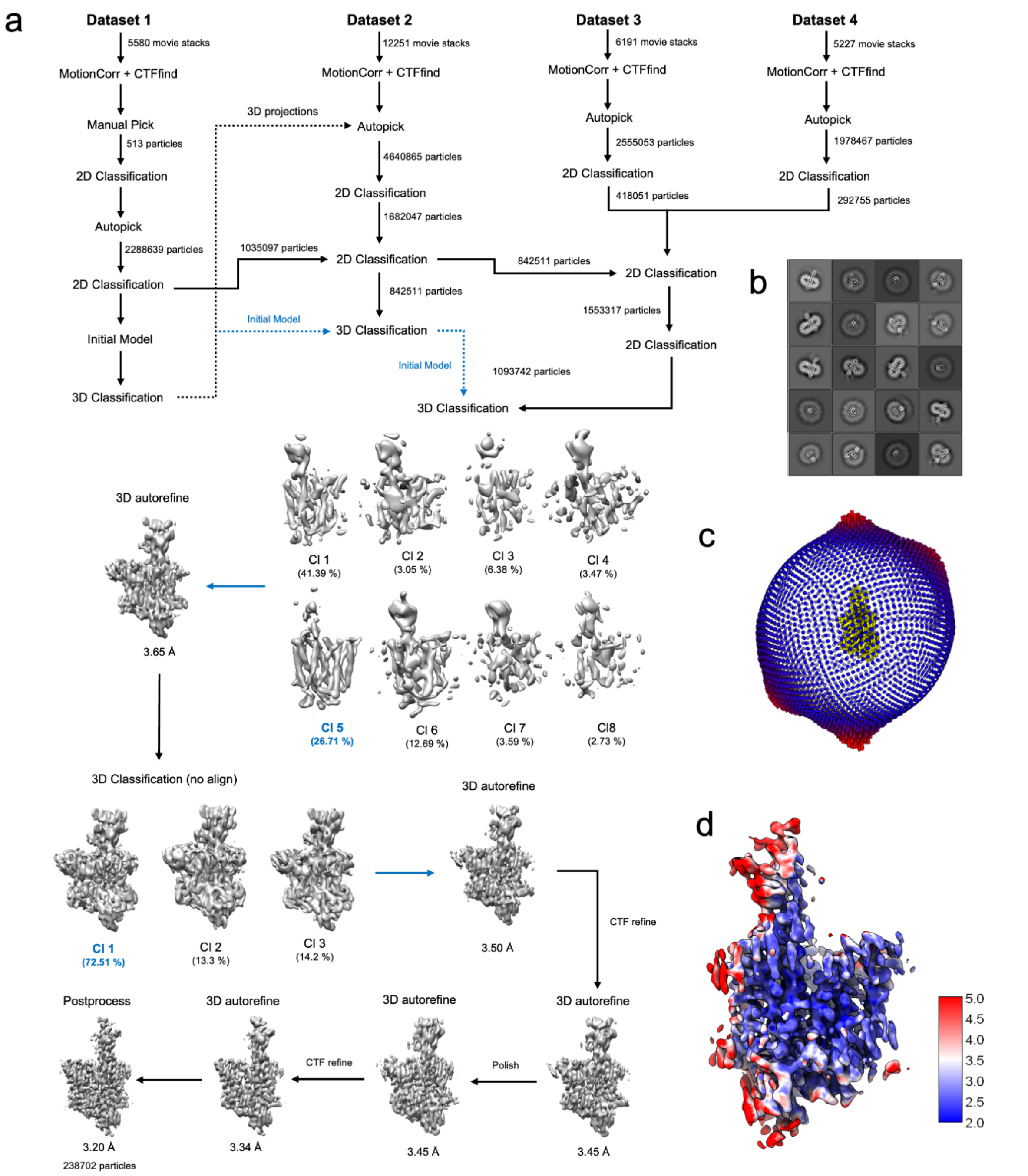


**Supplementary Fig. 2. Cryo-EM analysis of the apo-OCT3. a,** Cryo-EM image processing scheme used for 3D reconstruction of apo-OCT3. **b,** Representative 2D classes. **c,** Angular distribution histogram of the refined apo-OCT3 map. **d,** Local resolution map of apo-OCT3.

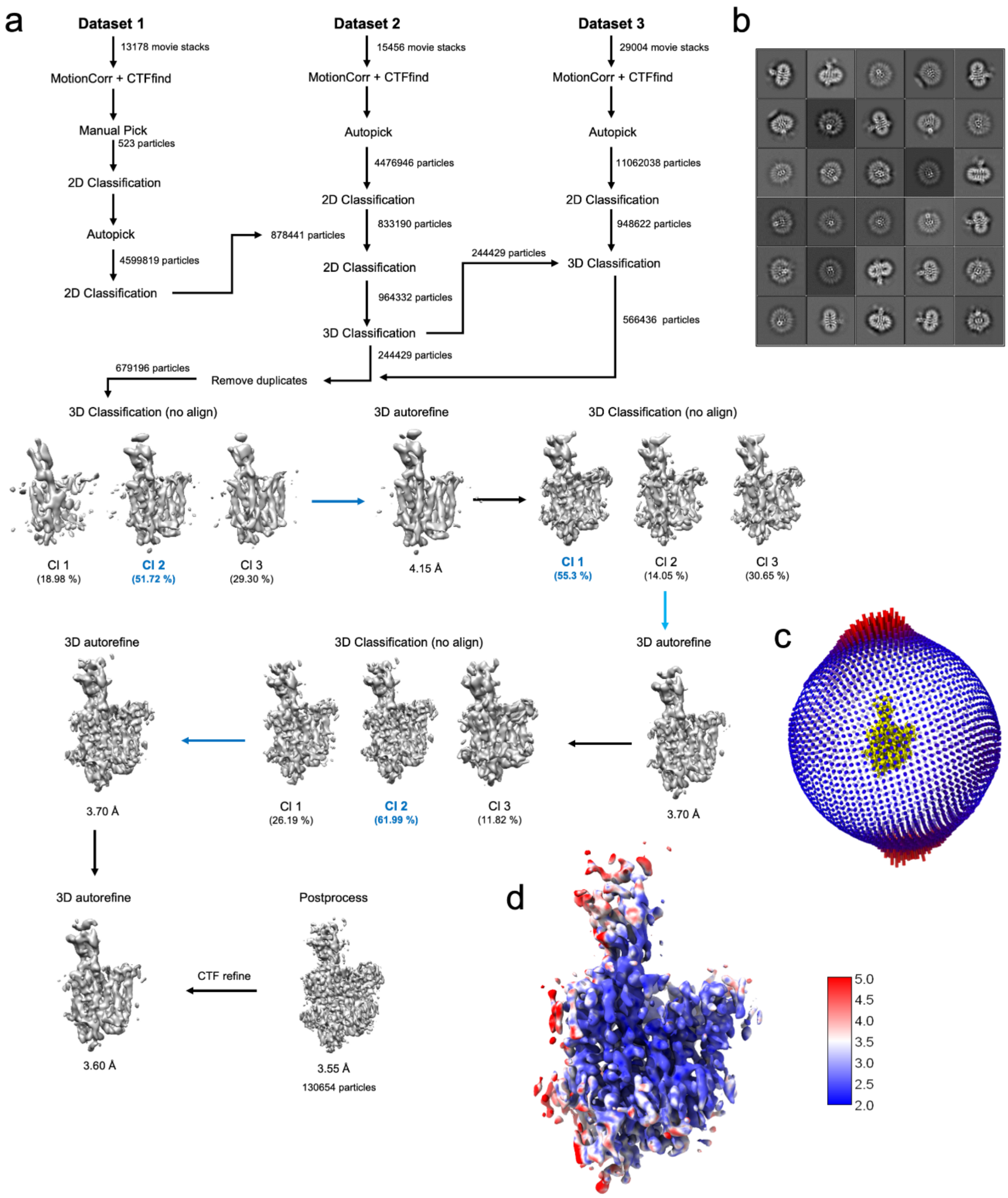


**Supplementary Fig. 3. Cryo-EM analysis of the OCT3-D22 complex. a,** Cryo-EM image processing scheme used for 3D reconstruction of OCT3-D22. **b,** Representative 2D classes. **c,** Angular distribution histogram of the refined apo-OCT3 map. **d,** Local resolution map of OCT3-D22.

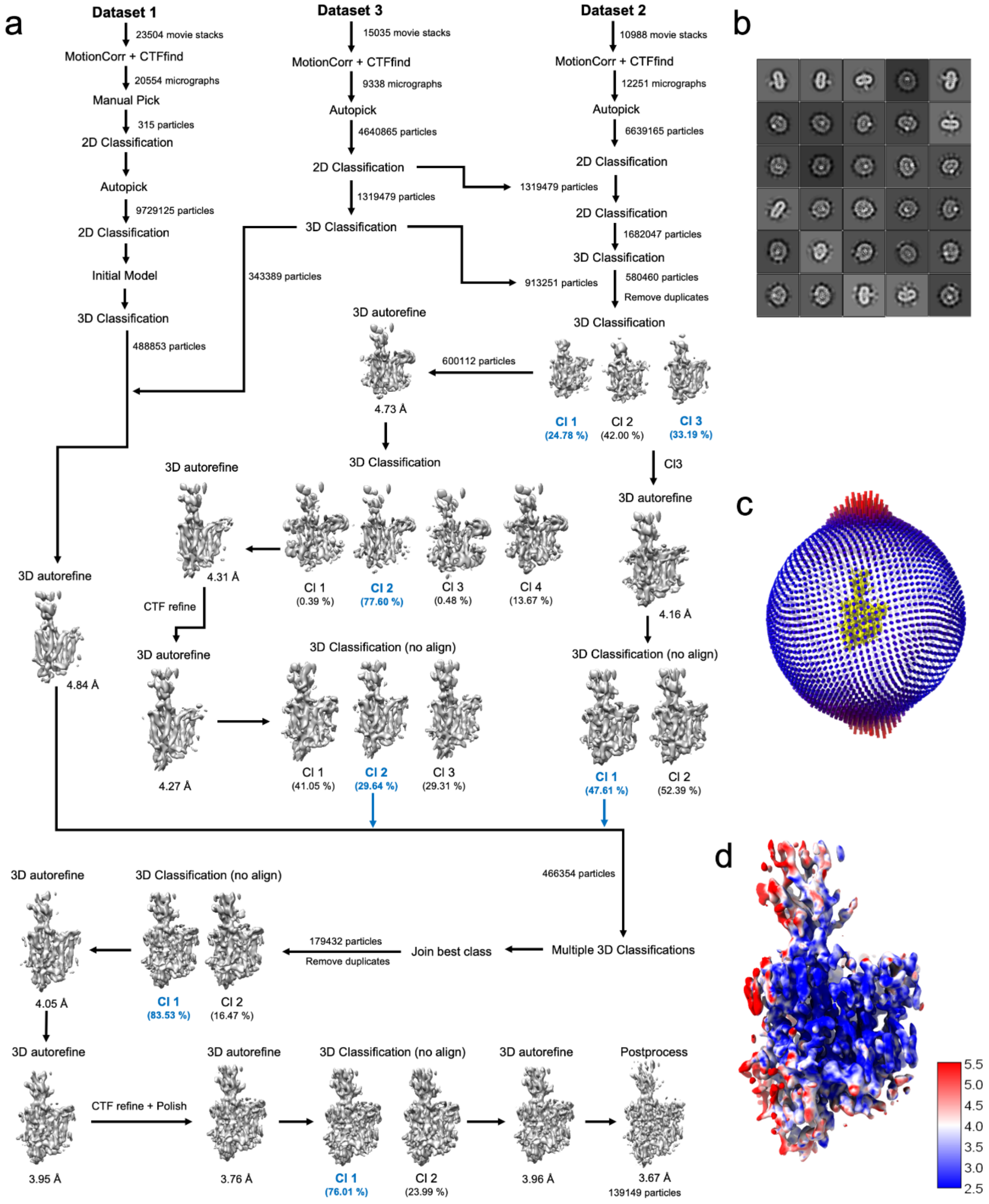


**Supplementary Fig. 4. Cryo-EM analysis of the OCT3-CORT complex. a,** Cryo-EM image processing scheme used for 3D reconstruction of OCT3-CORT. **b,** Representative 2D classes. **c,** Angular distribution histogram of the refined OCT3-CORT map. **d,** Local resolution map of OCT3-CORT.

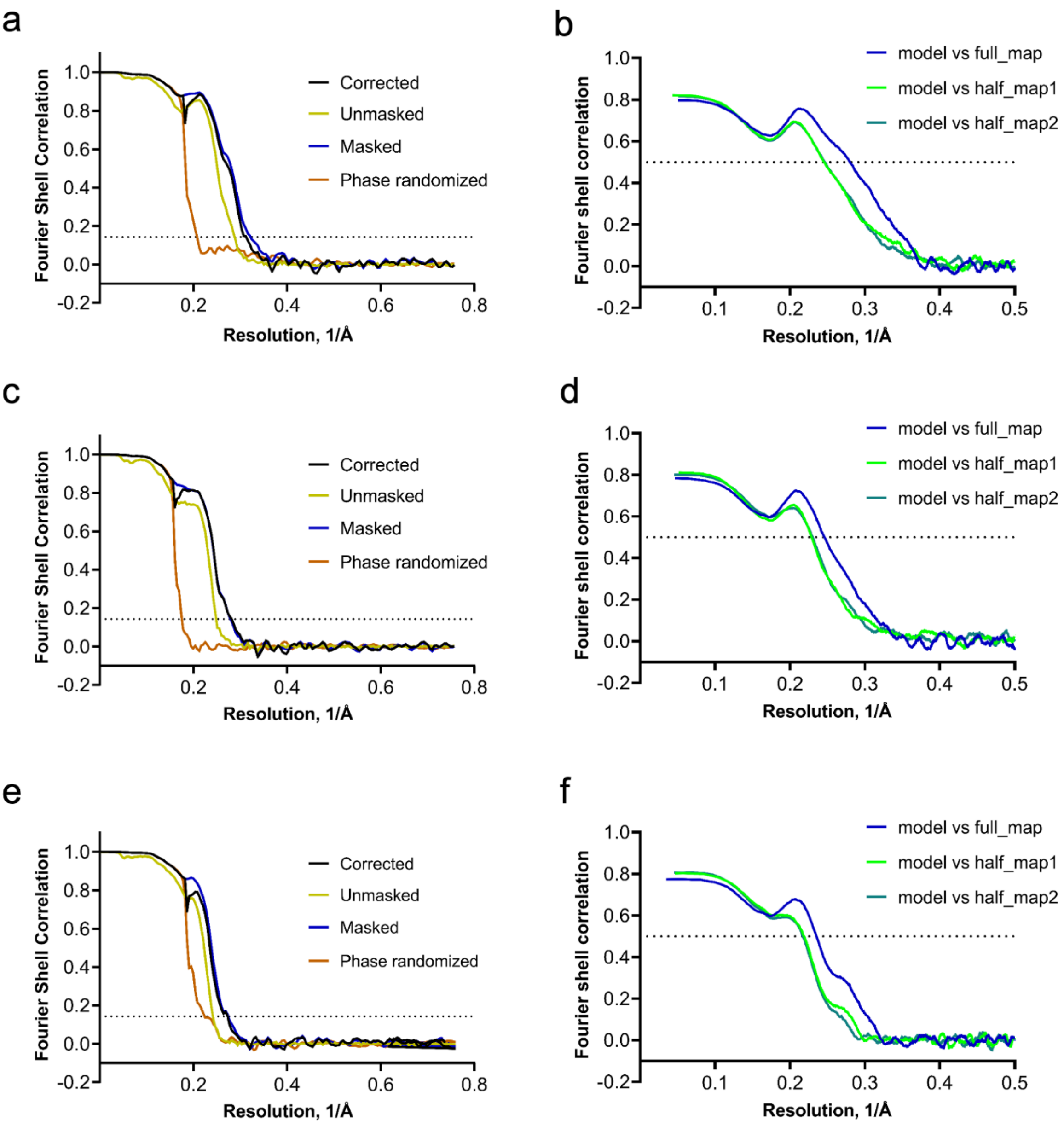


**Supplementary Fig. 5. Fourier shell correlation (FSC) curves of OCT3 cryo-EM maps. a,** FSC plot of the final 3D reconstruction and **b,** map to model FSC plot for apo-OCT3. **c,** FSC plot of the final 3D reconstruction and **d,** map to model FSC plot of OCT3-D22 complex. **e,** FSC plot of final 3D reconstruction and **f,** map to model FSC plot of OCT3-CORT complex.

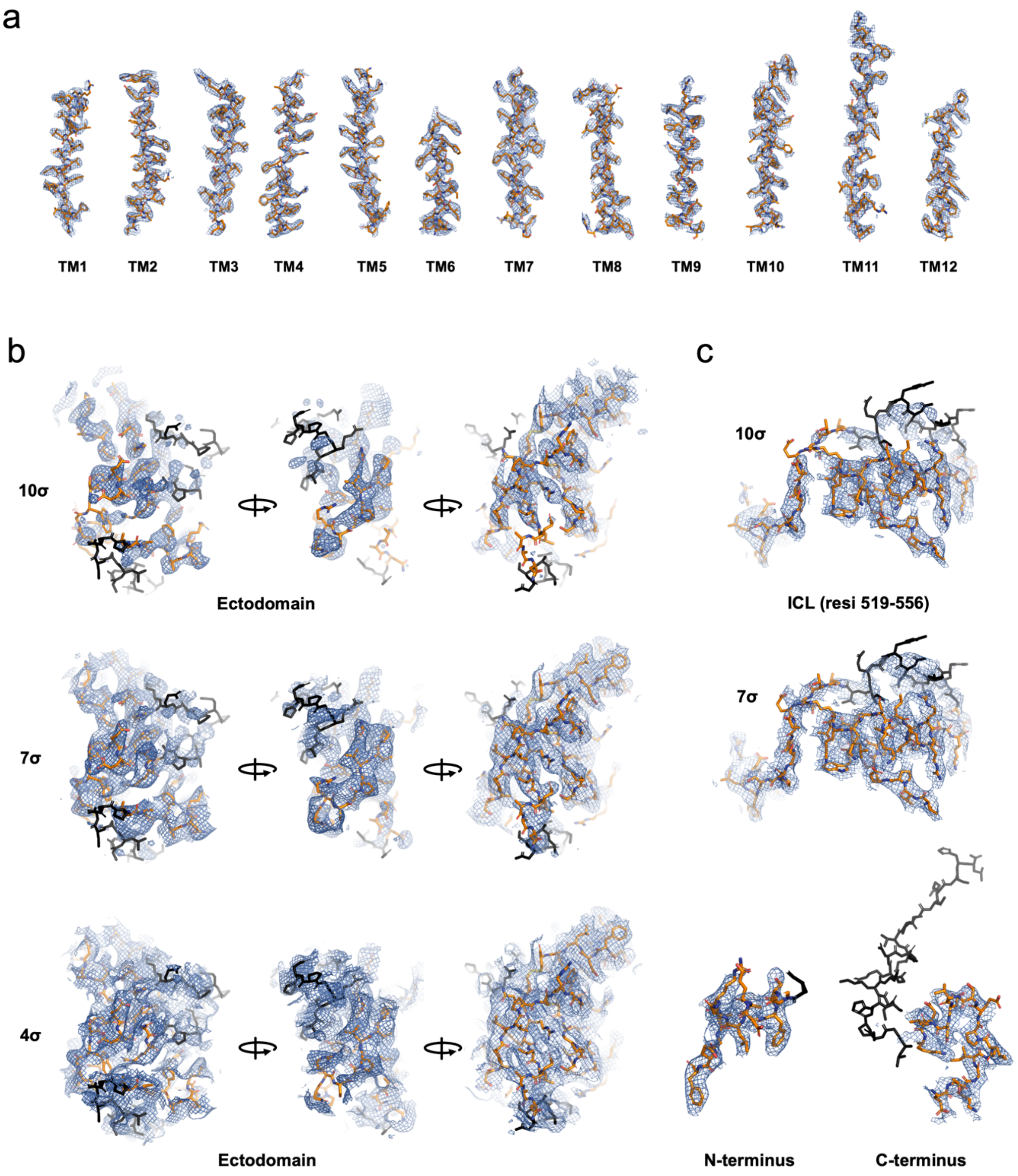


**Supplementary Fig. 6. Cryo-EM map features and model building of hOCT3. a,** Isolated density map for 12 TM helices of apo-OCT3 contoured at 12σ threshold level. **b,** Cryo-EM density features of ectodomain (ECD) contoured at 10σ, 7σ and 4σ levels. **c,** Cryo-EM density features for intracellular regions/domains including N- and C-terminus of OCT3. The residues modelled using AlphaFold are colored black.

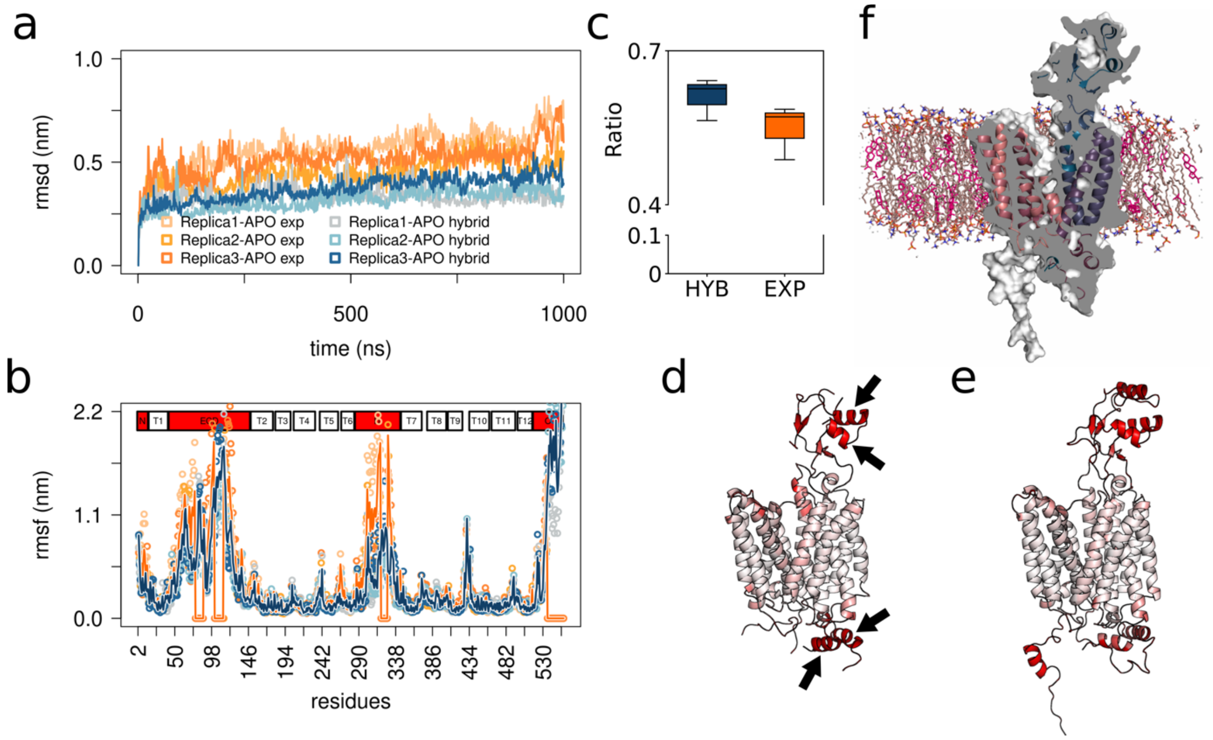


**Supplementary Fig. 7. Structural stability of hOCT3. a,** Time evolution of the root-mean-square deviation (RMSD) with respect to the respective starting structure. All resolved experimentally residues were included in the analysis (residue 2-74, 88-99, 114-316 and 328-531) for both the system containing only the experimentally determined coordinates (exp) as well as the hybrid system, which included residues predicted by AlphaFold (hybrid). Trajectories were fitted to the transmembrane helices (TMs) (residue TM1 (16-41), TM2 (149-178), TM3 (181-201), TM4 (205-233), TM5 (239-263), TM6 (267-284), TM7 (345-372), TM8 (379-404), TM9 (406-425), TM10 (434-460), TM11 (463-494) and TM12 (497-516)) of the respective starting structure before analysis. **b,** Per residue root-mean-square fluctuation (RMSF). Circles represent RMSF values of the independent system, colored as in panel a). The dark orange and blue lines show the averaged RMSF values for the exp and the hybrid system, respectively. Boxes at the top indicate the TMs (T1-T12), the extracellular domain (ECD), the intracellular loop region (I1) and the N- and C-termini (N & C). **c,** Box plot quantifying the average secondary structure content (calculated by using the dssp algorithm) of the helices the ECD (residue 75-87 and 100-113) and intracellular loop region I1 (residue 317-325 and 535-556), which are in contact with the AlphaFold predicted region. These are indicated by arrows in panel d. The averaged RMSF per residue are shown as β-factors for the simulation of the **d,** exp and **e,** OCT3 models. **f,** Slice through the membrane embedded hybrid model of OCT3, which is colored according to residue number. The dissected surface is represented in dark gray. The POPC and cholesterol molecules of the membrane are colored in brown and magenta, respectively.

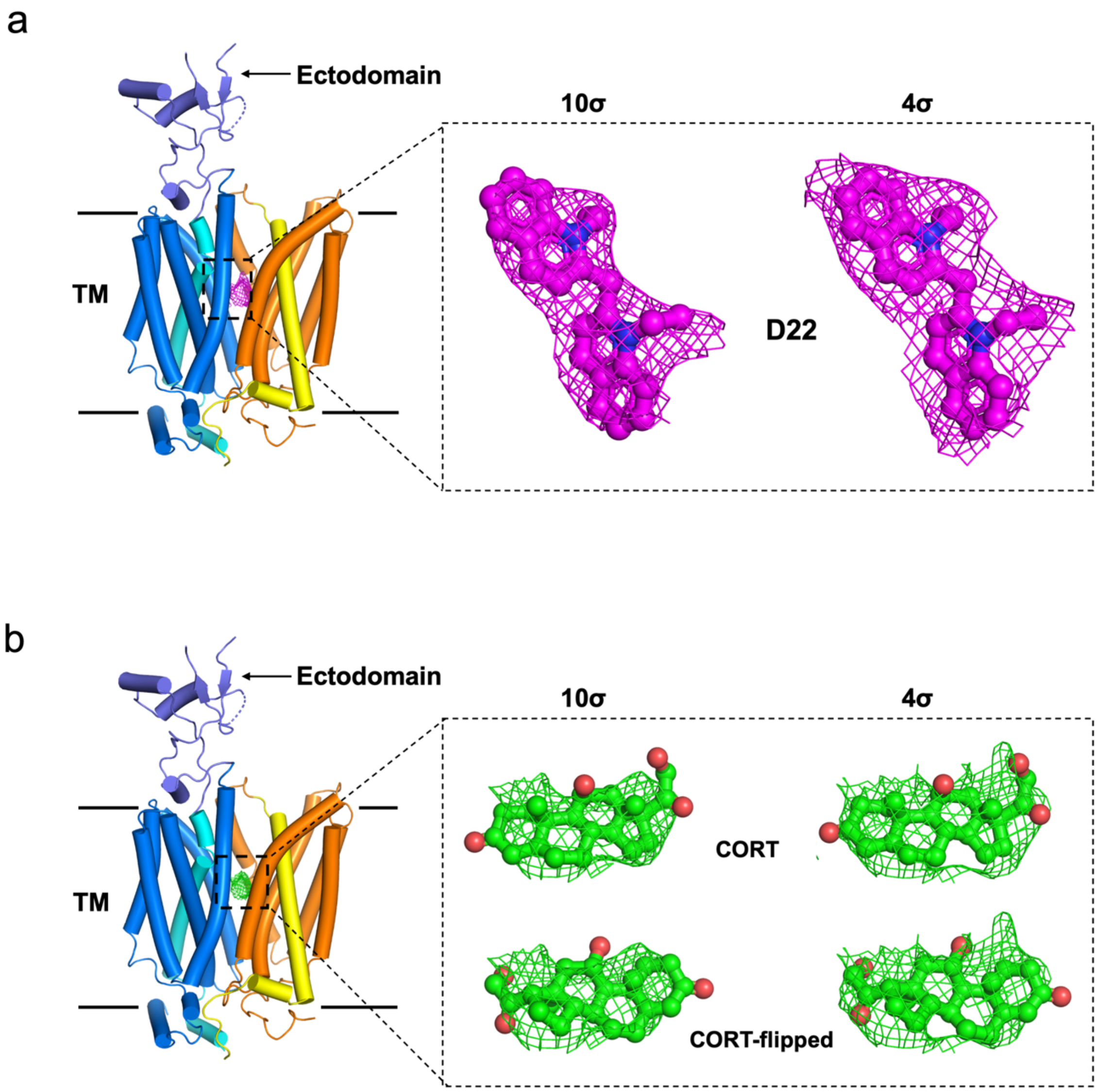


**Supplementary Fig. 8. Modelling of D22 and CORT in density maps. a,** Cartoon representation of OCT3 showing density of D22 (magenta mesh) at 10σ and 4 σ contour levels. **b,** Cartoon representation of OCT3 showing density of CORT (green mesh) occupying the center of the substrate translocation pathway. The CORT molecule was modelled in two alternate conformations labelled as CORT and CORT-flipped (~180 flipped orientations of CORT).

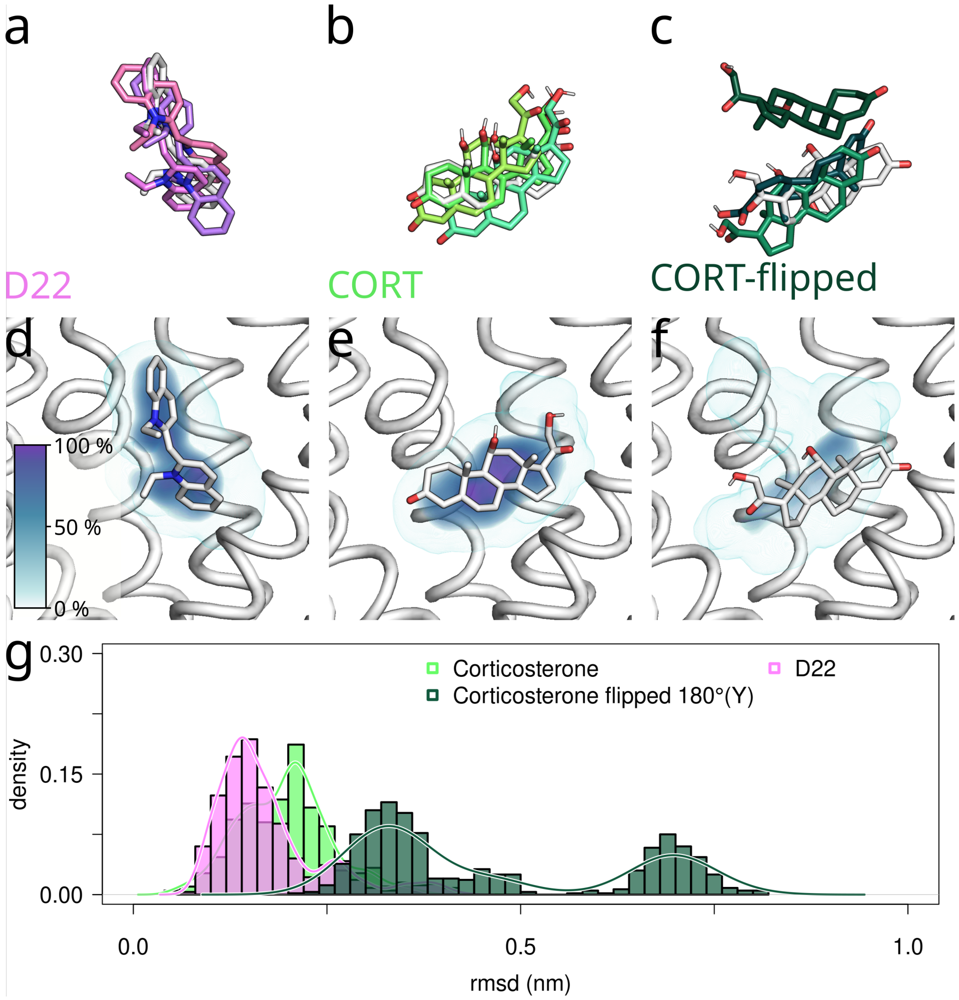


**Supplementary Fig. 9. Structural stability of ligands in their proposed binding site.** As it was difficult to correctly place CORT into the experimental density, the two possibilities labelled as CORT and CORT-flipped were simulated. The starting poses of **a,** D22 **b,** CORT and **c,** CORT-flipped are shown in white and the respective end poses at 150 ns are overlayed, after fitting all trajectories to the transmembrane helices. OCT3 is not shown for clarity. The occupancy in space of **d,** D22 **e,** CORT and **f,** CORT-flipped, averaged over the 3 trajectories (3 x 150 ns), are represented as volume density and colored according to the color legend. For reference, the OCT3 is shown in white together with the starting conformation of the ligand, while the OCT3 helices in front of the ligand are not shown. **h,** Histogram of RMSD values of the ligand with respect to the starting structure, after fitting the simulations to the transmembrane helices. The large RMSD of the flipped CORT allowed ruling out the second model of CORT placement.

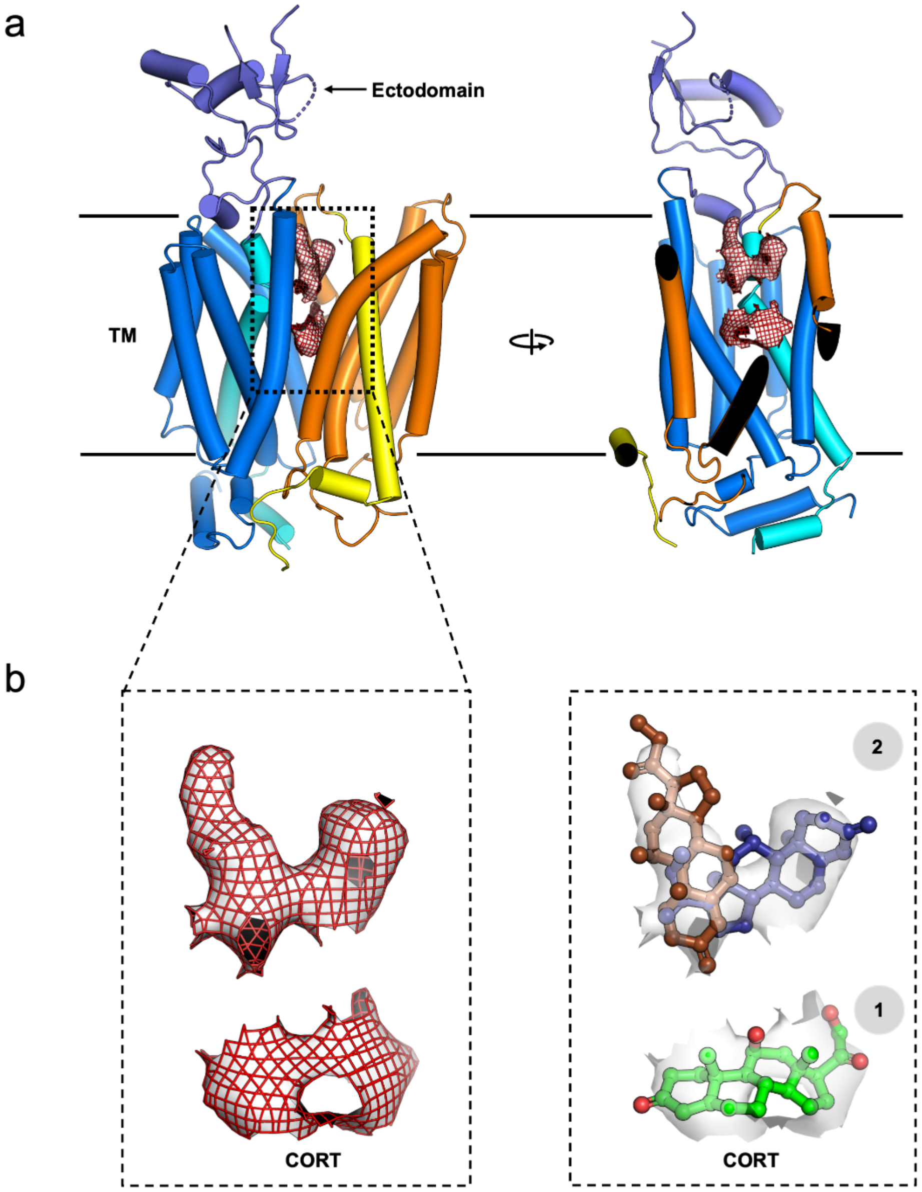


**Supplementary Fig. 10. Additional CORT densities above the substrate binding pocket of OCT3-CORT. a,** Cartoon representation of hOCT3 showing additional weak corticosterone or sterol-like density (shown as surface & mesh) at the entrance of the substrate translocation pathway. **b,** Electron density of sterol-like molecule contoured (4σ) threshold levels and potential alternate conformations of second CORT molecule are shown in brown and dark blue colored sticks.

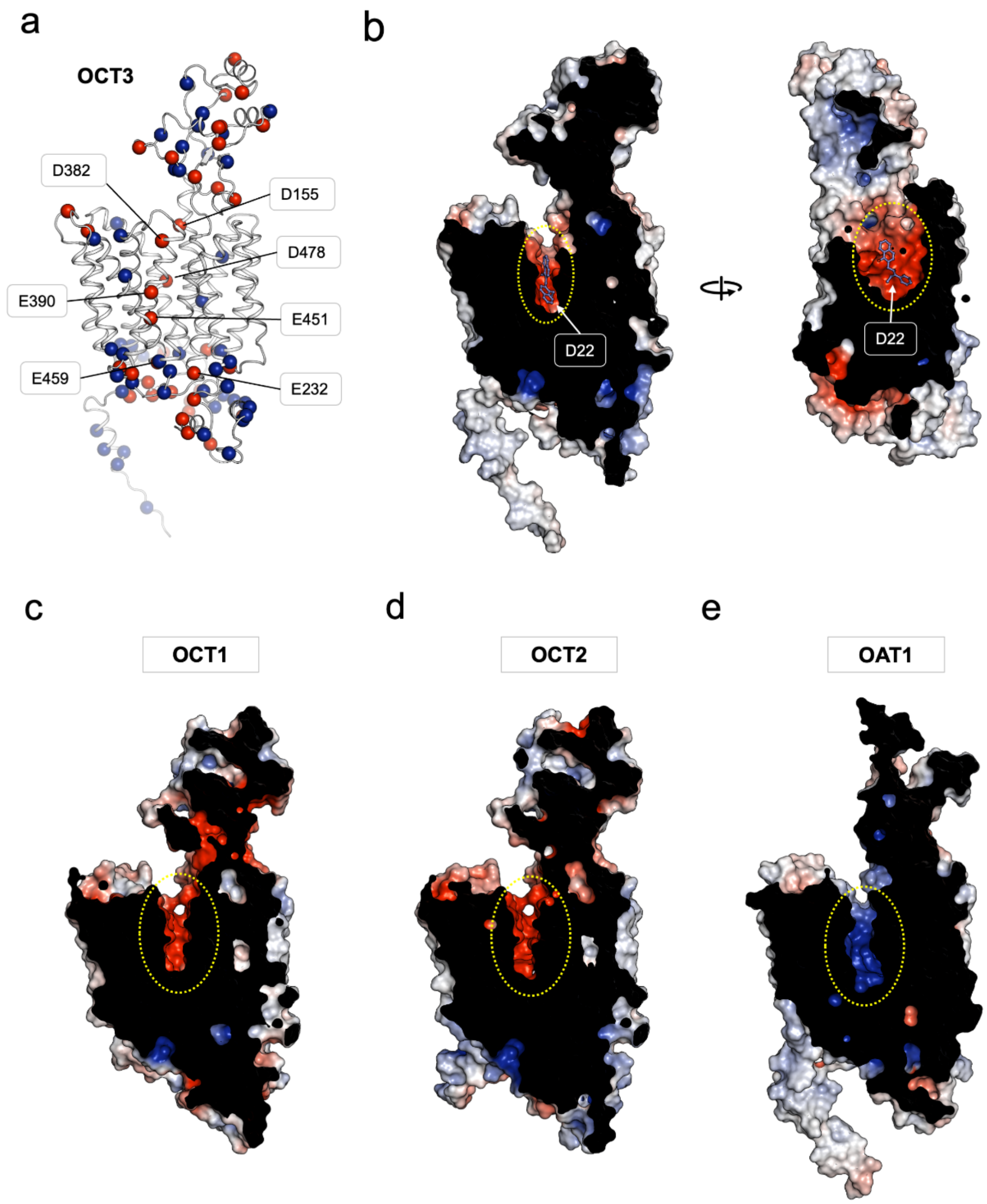


**Supplementary Fig. 11. Substrate binding pockets in OCTs and OATs. a,** Cartoon representation of the human OCT3 hybrid model depicting the distribution of positively (blue spheres) and negatively charged (red spheres) residues. **b,** The surface of OCT3-D22 coloured according to the calculated electrostatic potential, highlighting the negatively charged substrate binding site. **c-e,** Same as B for the homology model of the human OCT1, OCT2 and OAT1, indicating a charged substrate binding pockets. The OCT1, OCT2 and OAT3 homology were generated using SwissModel[10] with OCT3 as template. The sequence identity and modelling scores are tabulated in Supplementary Table 3. Electrostatic potential for each of the models was calculated with the Adaptive Poisson-Boltzmann Solver (APBS) module in Pymol and depicted on the same scale (-10 kT/e red, +10 kT/e blue).

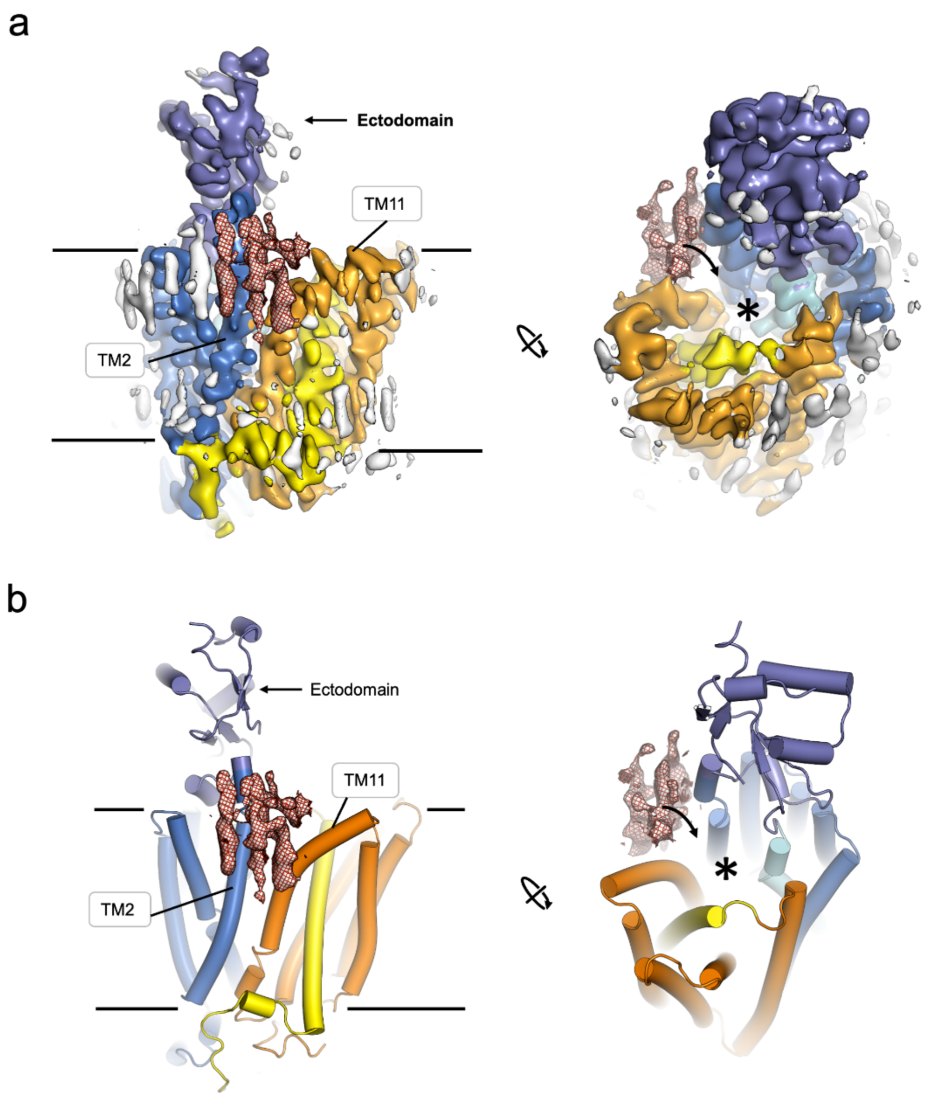


**Supplementary Fig. 12. Lateral lipid bilayer access to the substrate translocation pathway. a,** The cryo-EM map **b,** model of apo-OCT3 reveals densities that correspond to ordered lipid molecules in close proximity to a V-shaped opening formed by TM2 and TM11, separating the outer leaflet of the lipid bilayer from the substrate translocation pathway of OCT3.

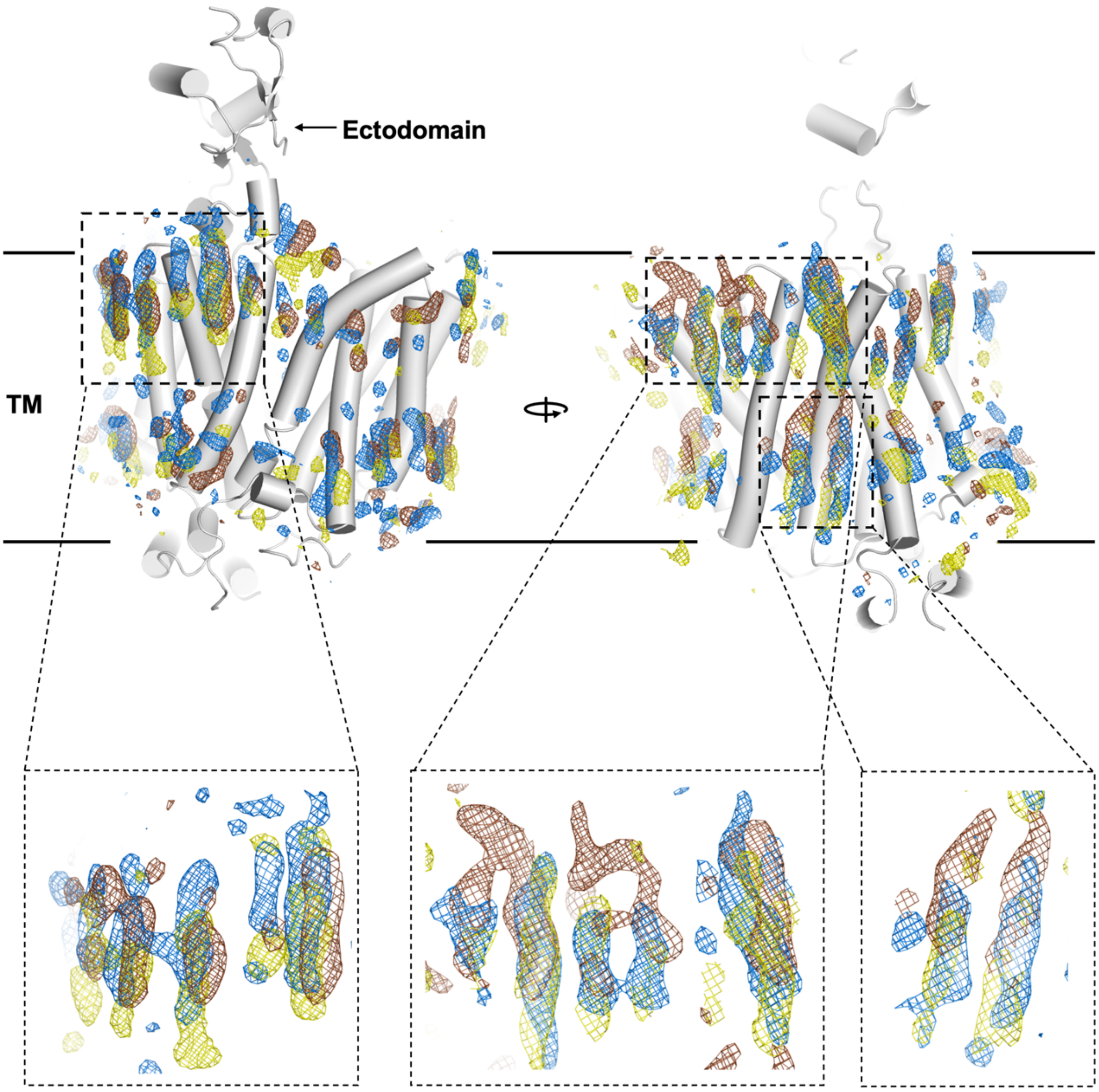


**Supplementary Fig. 13. Comparison of ordered lipid densities in OCT3 structures.** Cartoon representation of OCT3 model depicting ordered lipid-like densities on surface of OCT3 apo (marine), D22 (yellow) and CORT (red) bound states.

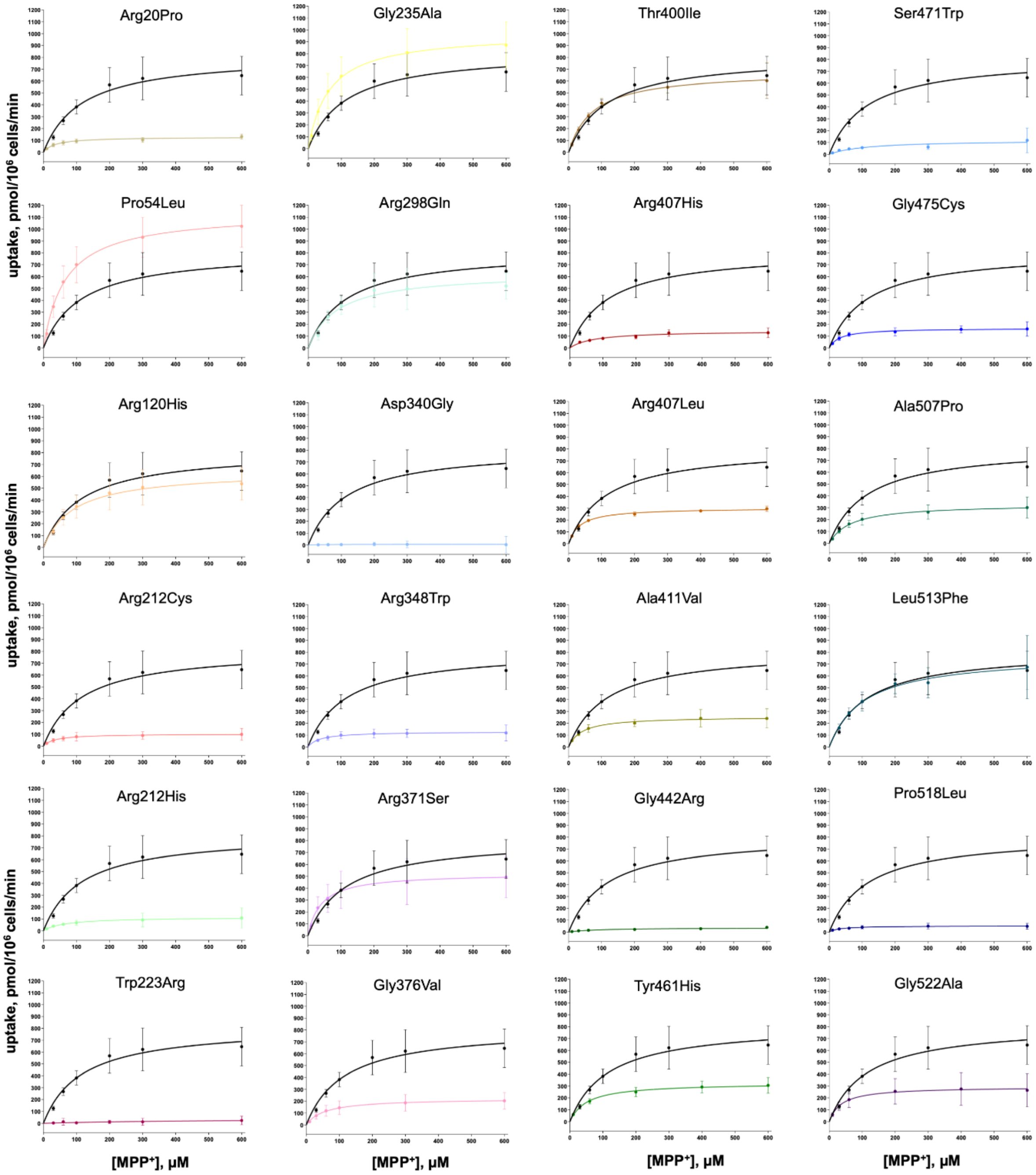


**Supplementary Fig. 14.** Uptake of $MPP^+$ by wildtype OCT3 (in black) and genetic variants expressed in HEK293 cells (n = 3 biologically independent experiments, in triplicate).

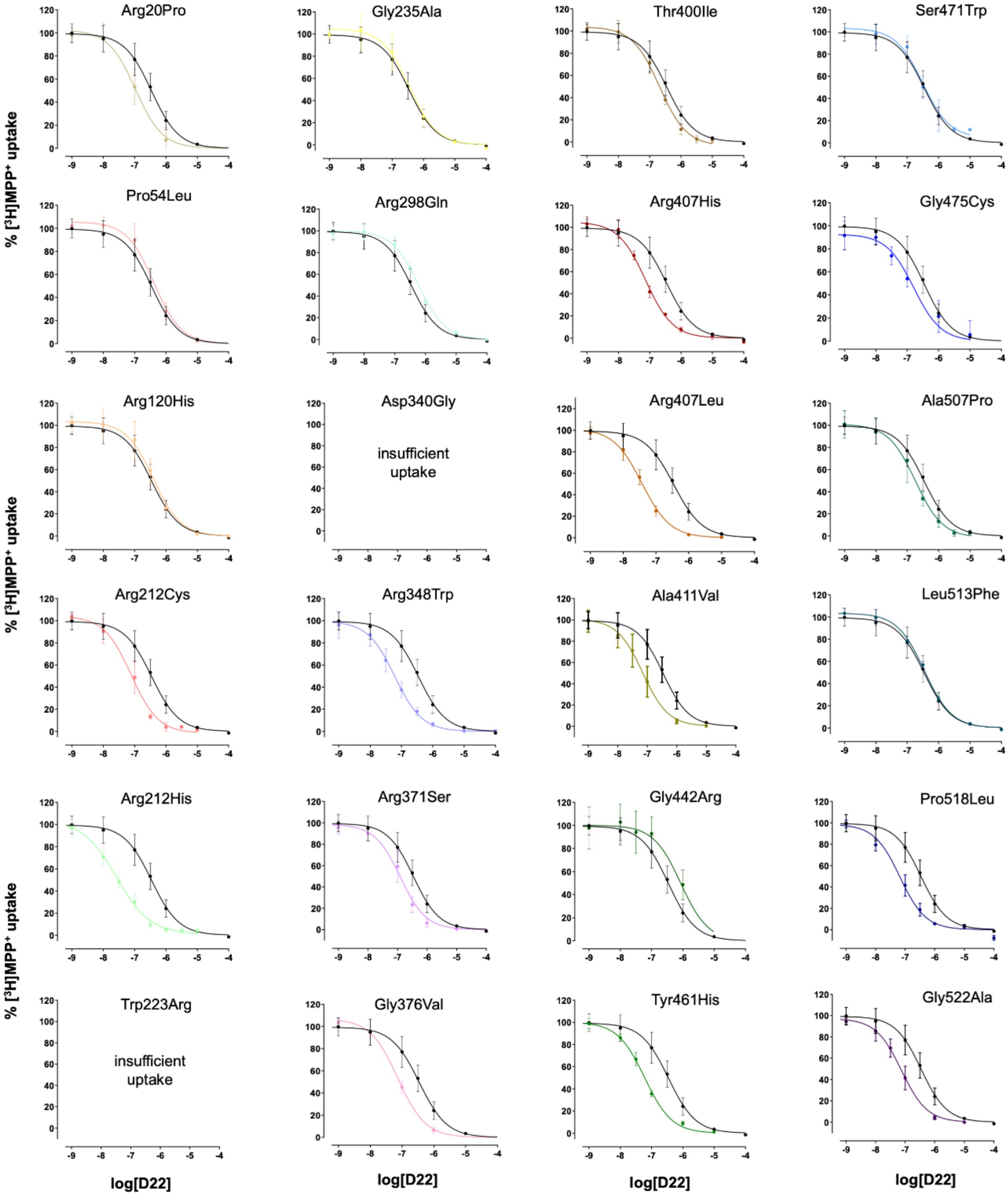


**Supplementary Fig. 15.** Inhibition of MPP$^+$ transport by wildtype hOCT3 (in black) and genetic variants expressed in HEK293 cells (n = 3 biologically independent experiments, in triplicate).

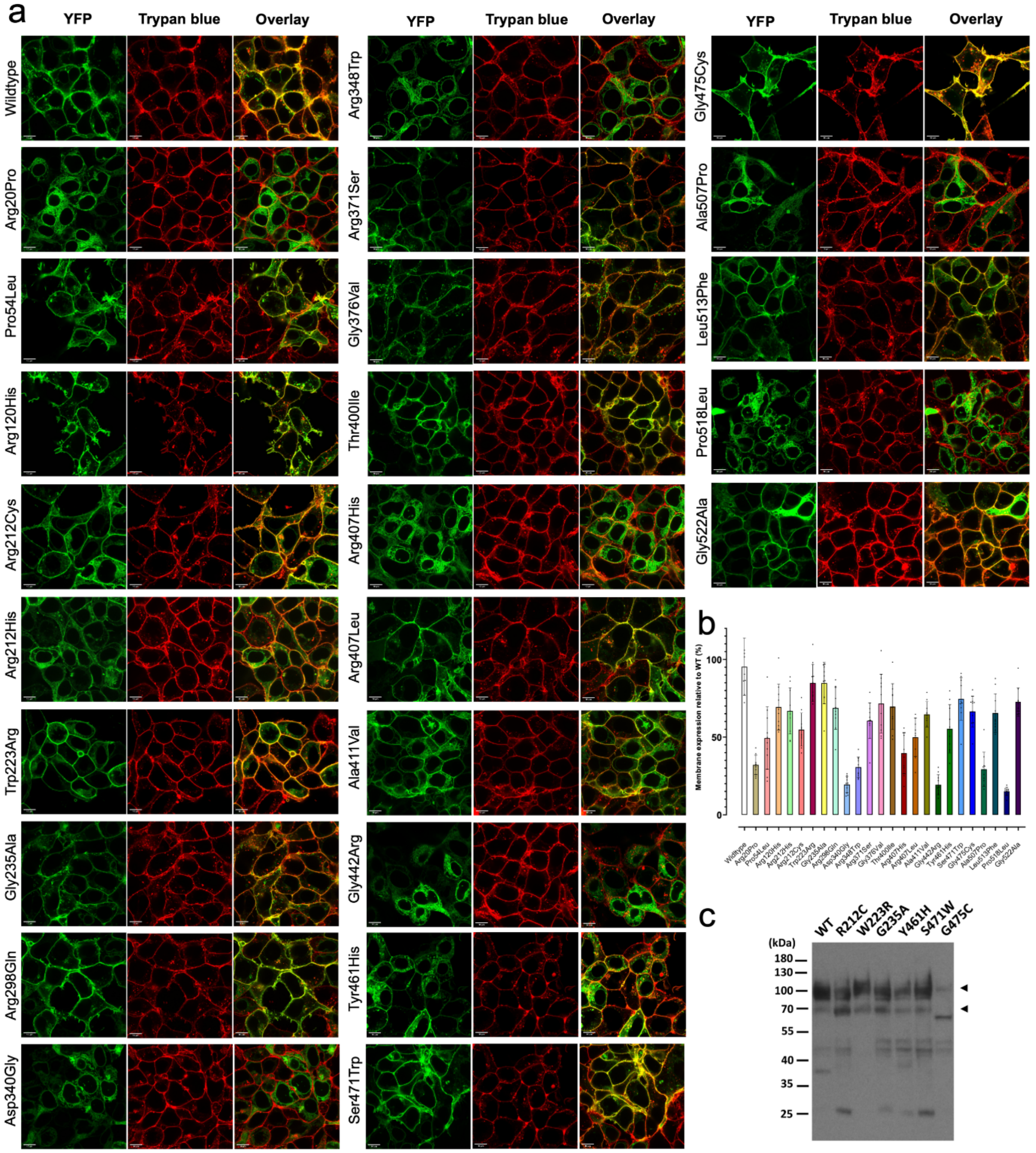


**Supplementary Fig. 16. a,** Representative confocal microscopy images of wildtype OCT3 and genetic variants expressed in HEK293 cells. Transporter was tagged with YFP and membrane stained with trypan blue. **b,** Analysis of surface expression of genetic variants, normalized to wildtype. Images were taken on three separate days and cell surface expression of cells measured in ImageJ. **c,** Immunoblot analysis performed against lysates from HEK293 cells expressing YFP-tagged OCT3 wildtype and mutants R212C, W223R, G235A, Y461H, S471W and G475C. Two populations of YFP-tagged OCT3 proteins (arrowheads) were detected by using an anti-GFP antibody. The numbers refer to the positions of pre-stained molecular weight markers.

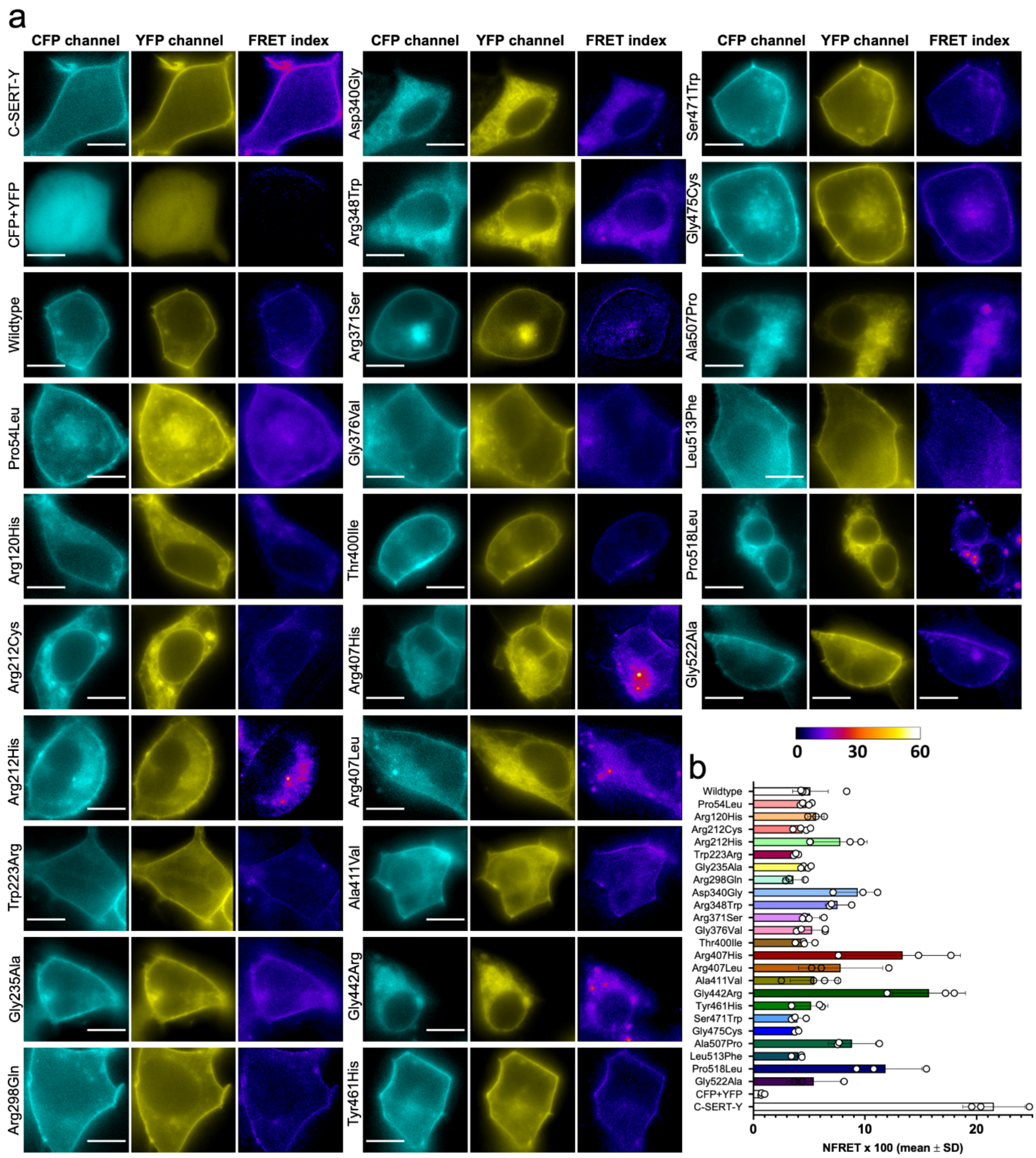


**Supplementary Fig. 17. a,** Representative Förster Resonance Energy Transfer (FRET) microscopy images of wildtype hOCT3 and genetic variants. N- and C-terminally tagged serotonin transporter (SERT) served as positive control and solely donor and acceptor fluorophore as negative. **b,** Analysis of NFRET values of analyzed proteins. Images were taken on three separate days and dots constitute averages per day.

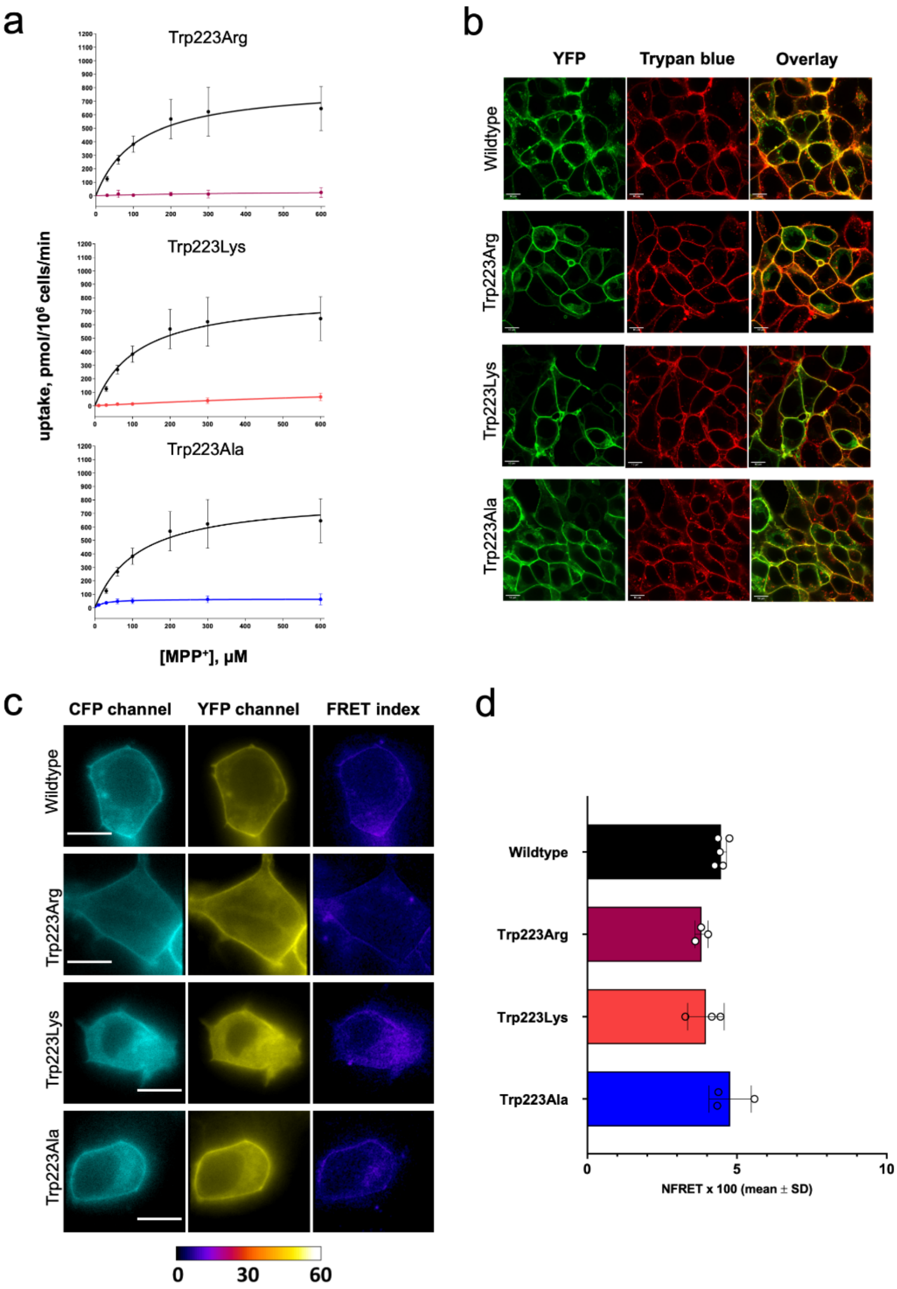


**Supplementary Fig. 18. a,** Uptake of $MPP^+$ by wildtype hOCT3 (in black), the genetic variant Trp223Arg and two variants not found in the genetic cohort: W223K and W223A (n=3 biologically independent experiments, in triplicate). **b,** Representative confocal microscopy images of wildtype OCT3 and variants expressed in HEK293 cells. Transporter was tagged with YFP and the membrane was stained with trypan blue. **c,** Representative Förster Resonance Energy Transfer (FRET) microscopy images of wildtype hOCT3 and genetic variants. **d,** Analysis of NFRET values of variants. Images were taken on three separate days and dots constitute averages per day.

**Supplementary Table 1.** Cryo-EM analysis and statistics

| Data collection | | | |
|---|---|---|---|
| | OCT3-apo | OCT3-D22 | OCT3-Corticosterone |
| Instrument | FEI Titan Krios / Gatan K3 Summit / | | |
| Magnification | 130000 | | |
| Voltage (kV) | 300 | | |
| Electron Dose (e-/Å2) | | | |
| Data-set 1 | 48 e-/Å$^2$ | 51 e-/Å$^2$ | 49 e-/Å$^2$ |
| Data-set 2 | 48 e-/Å$^2$ | 49 e-/Å$^2$ | 55 e-/Å$^2$ |
| Data-set 3 | 48 e-/Å$^2$ | 49 e-/Å$^2$ | 61 e-/Å$^2$ |
| Data-set 3 | 49 e-/Å$^2$ | - | - |
| Defocus range (μm) | -0.5 to -2.8 | -0.5 to -2.8 | -0.5 to -2.8 |
| Pixel size (Å) | 0.66 | 0.66 | 0.66 |
| **Refinement** | | | |
| Number of particles | 238702 | 130654 | 139149 |
| Map symmtery | C1 | C1 | C1 |
| Model resolution FSC threshold 0.5 | 3.20 | 3.55 | 3.67 |
| Map sharpening B-factor (Å) | -80 | -130 | -95 |
| Map CC | 0.75 | 0.72 | 0.73 |
| Model composition | | | |
| Protein residues/ligands | 495 | 487/1 | 478/1 |
| Bond length (r.m.s.d) | 0.004 | 0.004 | 0.014 |
| Bond angle (r.m.s.d) | 0.719 | 0.721 | 1.159 |
| **Validation** | | | |
| MolProbity score | 2.05 | 2.04 | 2.41 |
| Clashscore | 13.89 | 15.82 | 26.43 |
| Rotamer outlier (%) | 0.24 | 0.50 | 1.26 |
| Ramachandran plot | | | |
| Favoured (%) | 94.05 | 95.20 | 93.59 |
| Allowed (%) | 5.95 | 4.80 | 6.41 |
| Disallowed (%) | 0 | 0 | 0 |

**Supplementary Table 2.** pLDDT score of the hybrid OCT3 model generated with the help of AlphaFold.

| Residues range | Number of residues | Modelling method | pLDDT score (Alphafold) | | |
|---|---|---|---|---|---|
| | | | Min | Max | Average |
| 1-74 | 74 | EXP | 47.25 | 95.60 | 84.92 |
| **75-86** | **12** | **AF** | **39.82** | 79.19 | **52.57** |
| 87-99 | 13 | EXP | 56.85 | 86.22 | 77.70 |
| **100-113** | **14** | **AF** | **49.47** | 84.08 | **68.11** |
| 114-316 | 202 | EXP | 68.26 | 98.25 | 90.56 |
| **317-327** | **21** | **AF** | **53.93** | **86.17** | **67.45** |
| 328-535 | 207 | EXP | 37.85 | 97.71 | 85.91 |
| **536-556** | 21 | AF | 27.76 | 61.69 | 36.85 |

**Supplementary Table 3.** Statistics for the homology models of OCTs and OATs performed using Swiss model server and OCT3 as a template.

| **Name** | **Sequence identity (%)** | **Modelling score** | |
|---|---|---|---|
| | | **GMQE** | **QMEAN DisCo Global** |
| **OCT1** | 51.52 | 0.47 | 0.50 ± 0.05 |
| **OCT2** | 51.87 | 0.47 | 0.48 ± 0.05 |
| **OAT1** | 35.31 | 0.44 | 0.47 ± 0.05 |

**Supplementary Table 4:** Coding variants identified in SLC22A3 in the iPSYCH2012 cohort.

| Effect of mutation | Protein change | polyphen prediction | # controls carriers | #case carriers | Phentype(s) of cases | Allele frequncy in GnomAD |
|---|---|---|---|---|---|---|
| frameshift | Phe32fs | | <5 | <5 | ASD | ND |
| stop gained | Trp65* | | <5 | <5 | ASD | ND |
| frameshift | [illegible]u85fs (252delC) | | <5 | <5 | ASD, SZ | 3.20E-05 |
| frameshift | [illegible]u85fs (252dupC) | | 17 | 22 | ADHD,ASD,SZ,DP,BP | ND |
| frameshift | Gly393fs | | ND | <5 | ADHD | ND |
| splice donor | [illegible]1G>A splice donor | | <5 | <5 | ADHD, BP | ND |
| stop gained | Tyr185* | | <5 | ND | | ND |
| stop gained | Gln306* | | ND | <5 | ASD | 7.97E-06 |
| missense | Arg20Pro | D | <5 | ND | | ND |
| missense | Phe34Leu | B | <5 | <5 | ADHD, SZ | ND |
| missense | Pro54Leu | D | <5 | <5 | DP,BP | ND |
| missense | Gly80Asp | B | 5 | <5 | ASD | 6.55E-05 |
| missense | Pro83ser | B | <5 | <5 | ADHD, BP | 1.92E-05 |
| missense | Ala111Thr | B | ND | <5 | ADHD, SZ, DP | ND |
| missense | Pro126Arg | B | 35 | 69 | ADHD,ASD,SZ,DP,BP | 4.15E-05 |
| missense | Pro126Gln | B | ND | <5 | ADHD | 1.31E-03 |
| missense | Pro135ser | P | <5 | ND | | ND |
| missense | Val183Ile | B | <5 | ND | | 2.84E-05 |
| missense | Val194Ile | B | <5 | <5 | ADHD, ASD | 2.84E-05 |
| missense | Arg212Cys | D | <5 | 5 | ADHD, SZ, DP, BP | 3.55E-05 |
| missense | Arg212His | D | ND | <5 | SZ, DP | 2.13E-05 |
| missense | Trp223Arg | D | ND | <5 | ASD | ND |
| missense | Val230Ala | P | <5 | ND | | 3.98E-06 |
| missense | Gly235Ala | D | <5 | ND | | ND |
| missense | Ser236Leu | B | <5 | <5 | ASD | 4.38E-05 |
| missense | Thr 251Ser | P | <5 | ND | | ND |
| missense | Arg298Gln | P | 11 | 19 | ADHD,ASD,SZ,DP,BP | 4.78E-04 |
| missense | Arg310His | B | <5 | ND | | 2.83E-05 |
| missense | Leu319Phe | P | ND | <5 | ADHD, DP | ND |
| missense | Asp340Gly | D | ND | <5 | DP,BP | ND |
| missense | Val345Met | P | <5 | 7 | ADHD, ASD, DP, BP | ND |
| missense | Arg348Trp | D | ND | <5 | ADHD, ASD, SZ, BP | 8.14E-05 |
| missense | Arg371His | P | ND | <5 | ASD, SZ | 1.99E-05 |
| missense | Arg371Ser | D | ND | <5 | BP | 1.99E-05 |
| missense | Gly376Val | D | ND | <5 | SZ | ND |
| missense | Val388Met | B | <5 | 7 | ADHD, SZ, DP, BP | 1.20E-04 |
| missense | Leu391Val | P | ND | <5 | ASD | ND |
| missense | Thr400Ile | D | <5 | ND | | 2.79E-05 |
| missense | Ile401Val | P | ND | <5 | ASD, SZ | 1.99E-05 |
| missense | Arg407His | D | 21 | 70 | ADHD,ASD,SZ,DP,BP | 7.71E-04 |
| missense | Arg407Leu | D | ND | <5 | ADHD, ASD | ND |
| missense | Ala411Val | D | <5 | <5 | ADHD | 7.96E-06 |
| missense | LIle431Lys | P | 5 | 7 | ADHD, ASD, SZ | 9.20E-05 |
| missense | Arg435Ser | B | <5 | ND | | 2.39E-05 |
| missense | Gly442Arg | D | <5 | ND | | 3.98E-06 |
| missense | Tyr461His | D | <5 | ND | | ND |
| missense | Ser471Trp | D | ND | <5 | DP, BP | ND |
| missense | Gly475Cys | D | <5 | <5 | ADHD, ASD | 7.96E-06 |
| missense | Val494Met | P | ND | <5 | ADHD | 2.39E-05 |
| missense | Glu497Lys | P | ND | <5 | SZ | ND |
| missense | Ala507Pro | D | ND | <5 | ADHD | ND |
| missense | Leu513Phe | D | ND | <5 | ADHD | ND |
| missense | Pro518Leu | D | <5 | <5 | ASD | 2.83E-05 |
| missense | Gly522Ala | D | <5 | <5 | ASD | 2.83E-05 |
| missense | Val532Ala | B | ND | <5 | ADHD | ND |
| missense | Gly536Arg | B | ND | <5 | ASD | ND |
| missense | Pro538Ser | B | ND | <5 | ASD | 9.20E-06 |
| missense | Arg553His | P | <5 | <5 | ADHD | 3.57E-05 |

**Supplementary Table 5:** Coding variants identified in SLC22A3 in the iPSYCH2012 cohort. Carrier-based association analysis of coding SLC22A3 variants. Exome sequencing information on coding variants in SLC22A3 from the first phase of the iPSYCH case-cohort covering five major neuropsychiatric diagnosis: ADHD, ASD, schizophrenia, depression, and bipolar disorder covered by the ICD10 diagnoses: F90.0, F84.0, F84.0, F84.5, F84.8, F84. or F84.9, F20, F32-33, F30-F312. Diagnoses were obtained by linkage to the Danish Central Research Register, which contained information about all contacts up until 2016 t the time of linkage. Abbreviations: ADHD, attention-deficit/hyperactivity disorder; ASD, autism spectrum disorder; ICD10, International Classification of Diseases, 10th revision; iPSYCH, Integrative Psychiatric Research; LoF, Loss-of-function. Association analysis were performed using two-sided Fisher's exact test, *P<0.05, **P<0.001.

| | Carriers Controls (n=4885) | | Carriers Cases (n=12454) | | P-value (Two-sided Fischer's exact test) | OR [95% CI] |
|---|---|---|---|---|---|---|
| | N | % | N | % | | |
| Any coding SLC22A3 variants | 135 | 2.76 | 267 | 2.14 | ***P=0.0159** | **0.77 [0.624-0.949]** |
| LoF SLC22A3 variants | 25 | 0.512 | 29 | 0.233 | ****P=0.0057** | **0.45 [0.268-0.788]** |
| Missense SLC22A3 variants | 110 | 2.25 | 238 | 1.91 | P=0.149 | OR=0.846 [0.846-1.062] |

**Supplementary Table 6:** Pharmacological properties of wildtype OCT3 and genetic variants.

| hOCT3 isoform | $V_{max}$ (95%-CI) | $K_m$ (95%-CI) | $IC_{50}$ decynium-22 (µM, 95%-CI) | $IC_{50}$ corticosterone (µM, 95%-CI) |
|---|---|---|---|---|
| Wildtype | 819.8 (696.2-943.4) | 114.5 (65.73-163.4) | 0.35 (0.27-0.46) | 1.33 (1.17-1.50) |
| Arg20Pro | 129.0 (117.9-140.0) | 35.06 (23.46-46.66) | 0.11 (0.08-0.15) | - |
| Pro54Leu | 1142.0 (1041.0-1243.0) | 66.21 (47.28-85.13) | 0.41 (0.32-0.53) | - |
| Arg120His | 644.2 (546.3-742.1) | 91.52 (48.94-134.1) | 0.41 (0.32-0.52) | - |
| Arg212Cys | 103.3 (85.4-121.2) | 33.72 (11.18-56.25) | 0.08 (0.07-0.11) | 0.12 (0.11-0.14) |
| Arg212His | 115.5 (82.50-148.5) | 65.21 (4.13-126.3) | 0.03 (0.02-0.03) | - |
| Trp223Arg | - | - | insufficient uptake | |
| Gly235Ala | 972.50 (861.7-1083.0) | 62.38 (39.01-85.76) | 0.38 (0.26-0.57) | 0.91 (0.80-1.05) |
| Arg298Gln | 634.2 (537.3-731.1) | 86.65 (45.33-128.0) | 0.57 (0.49-0.65) | - |
| Asp340Gly | - | - | insufficient uptake | |
| Arg348Trp | 129.2 (107.5-150.9) | 35.86 (9.28-62.43) | 0.06 (0.05-0.08) | - |
| Arg371Ser | 524.1 (456.5-591.7) | 39.4 (20.19-58.61) | 0.12 (0.10-0.15) | - |
| Gly376Val | 221.9 (186.6-257.3) | 54.87 (25.15-84.59) | 0.07 (0.06-0.08) | - |
| Thr400Ile | 682.9 (629.7-736.2) | 72.94 (54.98-90.89) | 0.21 (0.17-0.25) | - |
| Arg407His | 144.3 (126.3-162.3) | 74.29 (43.74-104.8) | 0.07 (0.06-0.08) | - |
| Arg407Leu | 302.6 (289.4-315.8) | 33.51 (27.18-39.84) | 0.04 (0.03-0.04) | - |
| Ala411Val | 256.3 (229.3-283.3) | 40.25 (22..67-57.83) | 0.07 (0.06-0.09) | - |
| Gly442Arg | 36.24 (30.86-41.62) | 90.87 (51.59-130.1) | 0.85 (0.55-1.32) | - |
| Tyr461His | 325.8 (306.9-344.7) | 51.11 (39.36-62.86) | 0.06 (0.05-0.06) | 0.08 (0.06-0.11) |
| Ser471Trp | 119.4 (75.11-163.8) | 111.8 (0.0-229.3) | 0.16 (0.10 0.27) | 0.66 (0.43-1.01) |
| Gly475Cys | 165.3 (148.8-181.8) | 31.54 (18.29-44.80) | 0.16 (0.12-0.22) | 0.86 (0.71-1.04) |
| Ala507Pro | 330.3 (293.1-367.6) | 65.64 (41.64-89.63) | 0.18 (0.13-0.26) | - |
| Leu513Phe | 780.2 (654.6-905.9) | 106.0 (56.41-155.6) | 0.36 (0.32-0.41) | - |
| Pro518Leu | 52.01 (42.74-61.28) | 30.06 (7.53-52.59) | 0.07 (0.06-0.08) | - |
| Gly522Ala | 293.4 (241.2-345.7) | 36.51 (9.71-63.31) | 0.07 (0.06-0.89) | - |
